# The transcriptionally permissive chromatin state of ES cells is acutely tuned to translational output

**DOI:** 10.1101/167239

**Authors:** Aydan Bulut-Karslioglu, Trisha A. Macrae, Juan A. Oses-Prieto, Sergio Covarrubias, Michelle Percharde, Gregory Ku, Aaron Diaz, Michael T. McManus, Alma L. Burlingame, Miguel Ramalho-Santos

## Abstract

A permissive chromatin environment coupled to hypertranscription is critical to drive the rapid proliferation of Embryonic Stem (ES) cells and peri-implantation embryos. We carried out a genome-wide screen to systematically dissect the regulation of the euchromatic state of ES cells. The results reveal that activity of cellular growth pathways, prominently protein synthesis, perpetuates the euchromatic state and hypertranscription of ES cells. Acute, mild inhibition of translation results in rapid depletion of euchromatic marks in ES cells and blastocysts, concurrent with delocalization of RNA polymerase II and reduction in nascent transcription. Remarkably, reduced translational output leads to rewiring of open chromatin within 3 hours, including decreased accessibility at a subset of active developmental enhancers and increased accessibility at histone genes and transposable elements. Using a proteome-scale analysis, we show that several euchromatin regulators are unstable proteins and thus continuously depend on a high translational output. We propose that this mechanistic interdependence of euchromatin, transcription and translation sets the pace of proliferation at peri-implantation and may be employed generally by stem/progenitor cells.

## INTRODUCTION

Stem and progenitor cells often display a distinct chromatin landscape associated with high levels of transcriptional activity (Gaspar-Maia et al., 2011; Percharde et al., 2017a; Reddien and Sánchez Alvarado, 2004; Spangrude et al., 1988). This chromatin state has been extensively studied in ES cells cultured in serum, which represent the rapidly proliferating pluripotent cells of the peri-implantation embryo (Smith, 2017). ES cells and pluripotent cells of the blastocyst display a remarkably decondensed chromatin pattern with low levels of compact heterochromatin (Ahmed et al., 2010; Efroni et al., 2008) and high levels of histone marks associated with transcriptional activity, such as H3/H4 acetylation and H3K4me3 (Ang et al., 2011; Krejcí et al., 2009; Lee et al., 2004). In agreement, ES cells are in a state of hypertranscription (Percharde et al., 2017a) that includes global elevation of nascent transcriptional output (Efroni et al., 2008).

Several factors have been implicated in the regulation of the permissive chromatin state of ES cells, including the histone acetyltransferases Tip60/p400 (Fazzio et al., 2008; Ravens et al., 2015) and Mof (X. Li et al., 2012), the trithorax group protein Ash2l (Wan et al., 2013) and the ATP-dependent chromatin remodelers Ino80 (Wang et al., 2014) and Chd1 (Gaspar-Maia et al., 2009; Guzman-Ayala et al., 2014). We have shown that Chd1 binds broadly to the transcribed portion of the genome and promotes hypertranscription by both RNA Pol I and II in ES cells (Gaspar-Maia et al., 2009; Guzman-Ayala et al., 2014). This Chd1-driven state of elevated transcription is essential for growth of pluripotent epiblast cells of the mouse embryo at the time of implantation (Guzman-Ayala et al., 2014) and of hematopoietic stem/progenitor cells emerging from the endothelium at mid-gestation (Koh et al., 2015). These data indicate that a permissive chromatin associated with global hypertranscription is required for developmental transitions that involve rapid proliferation of stem/progenitor cells. Despite these insights, the regulation of the permissive chromatin state of ES cells has not been dissected on a genome-wide scale. Moreover, a key question remains to be answered: how is hypertranscription set to the needs of rapidly proliferating pluripotent stem cells in the growing embryo? In other words, how do pluripotent stem cells sense when not enough or too much transcription is occurring, and adjust their chromatin state accordingly?

We report here a genome-wide RNAi screen to systematically probe the permissive chromatin state of ES cells. Integrated analyses at the functional, chromatin, transcriptional and proteome level reveal that the growth capacity of ES cells, specifically the translational output, directly promotes a permissive chromatin environment, at least in part by maintaining the levels of unstable euchromatin regulators. The results reveal a dynamic positive feedback loop between permissive chromatin and translation that drives proliferation of pluripotent cells and may be tuned by signaling and availability of nutrients.

## RESULTS

### A genome-wide RNAi screen identifies new regulators of euchromatin in ES cells

We sought to generate a live-cell reporter for euchromatin that would allow dissection of the dynamics and regulation of the euchromatic state of ES cells. The histone mark H3K4me3 is associated with active transcription, is directly and specifically bound by the double chromodomains of Chd1 (Flanagan et al., 2005) and is present at high levels in undifferentiated ES cells (Ang et al., 2011). We therefore generated mouse ES cells expressing a fusion between the Chd1 chromodomains and EGFP (referred to as Chd1chr-EGFP). As a control, we used ES cells expressing an Hp1α-EGFP fusion protein (Bulut-Karslioglu et al., 2014), which recognizes H3K9me3, a mark of constitutive heterochromatin. As anticipated, fluorescence in Chd1chr-EGFP ES cells is broadly distributed throughout the nucleus, whereas it is restricted to DAPI-dense heterochromatin in Hp1α-EGFP ES cells (Figure S1A).

To assess whether the Chd1chr-EGFP reporter responds to manipulations of the chromatin state, we first knocked down Wdr5, a core component of MLL complexes that deposit H3K4me3 (Ang et al., 2011) (Figure S1B). Wdr5 knock-down leads to a decrease in Chd1chr-EGFP reporter intensity within 3 days, whereas the Hp1α-EGFP reporter remains unaffected (Figure S1C). Similarly, we observed a rapid decrease in Chd1chr-EGFP intensity upon retinoic acid (RA)-mediated differentiation of ES cells for 2 days, prior to any detectable changes in the activity of the Hp1α-EGFP reporter (Figure S1D). Taken together, these results indicate that the Chd1chr-EGFP reporter responds as expected to perturbations of the euchromatic landscape of ES cells.

We next used the Chd1chr-EGFP ES cells to perform a genome-wide RNAi screen (Figure 1A). ES cells were infected with an ultra-complex lentiviral shRNA library comprised of ~30 shRNAs per gene. Cells were cultured for 3 days in serum/ leukemia inhibitory factor (LIF) medium and subsequently sorted into GFP^low^ and GFPhigh populations by flow cytometry. Integrated shRNAs were isolated, amplified and sequenced. Differential enrichments in shRNAs per gene recovered from the GFP^low^ and GFPhigh populations were used to estimate effect size, using HiTSelect (Diaz et al., 2015) (see Experimental Procedures for details). We previously used a similar approach to systematically identify barriers to human iPS cell generation (Qin et al., 2014). Applying this method at a 5% false discovery rate cutoff, we identified 486 genes whose knockdown associated with lower Chd1chr-EGFP fluorescence (Figure 1B and Table S1). These genes are thus putative positive regulators of the euchromatic state of ES cells. In agreement, this set of genes includes several previously described regulators of ES cell chromatin, including Chd1 itself (Gaspar-Maia et al., 2009), Hira (Banaszynski et al., 2013; Meshorer et al., 2006), Tip60/Kat5, p400 (Fazzio et al., 2008), Kmt2b (Denissov et al., 2014) and Ino80 (Wang et al., 2014).

**Figure 1.**
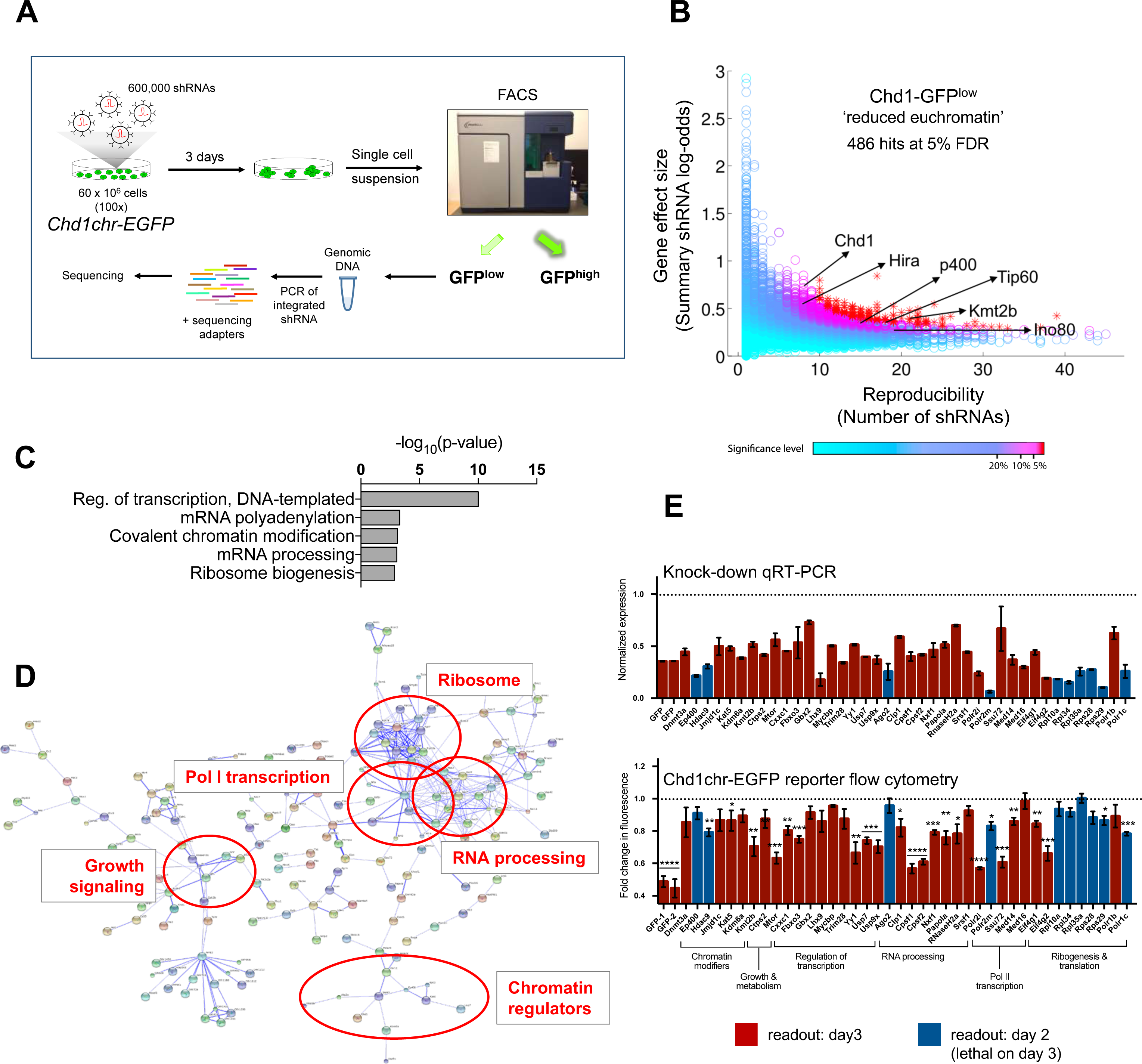
A genome-wide RNAi screen identifies regulators of euchromatin in ES cells. (A) RNAi screen workflow. (B) Results of the RNAi screen results for the genes with shRNAs enriched in the GFP^low^ fraction. Each circle annotates a gene tested in the screen. Published regulators of open chromatin in ES cells are indicated by arrows. See Table S1 for the full screen results. (C) Gene ontology (GO) terms associated with significant RNAi screen hits. See Table S2 for the full list of GO terms. (D) Protein interaction network of significant RNAi screen hits generated using STRING. (E) Secondary siRNA screen results. Genes were selected to represent each of the major pathways enriched in (C) and (D). Upper panel shows knockdown levels by qRT-PCR, normalized to scrambled siRNA control. Lower panel shows fluorescence level of the Chd1chr-EGFP reporter upon each knockdown. Readouts for both assays were measured on day 3 post-knockdown (red) or on day 2 (blue) if the knockdown was lethal by day 3. Error bars show mean ± standard deviation (SD) of 4 technical replicates. Graph is representative of 2 biological replicates. Statistical test performed is two-tailed t-test. *, **, ***, **** = p<0.05, 0.01, 0.001,0.0001.

We clustered screen hits according to functional annotations (Gene Ontology, GO) and protein interaction data (Figure 1C and D, Table S2). As expected, regulation of transcription and chromatin emerge as key processes modulating ES cell euchromatin. Intriguingly, many factors involved in cellular growth and protein synthesis, notably RNA Polymerase (Pol) I complex components, ribosomal proteins and translation factors, are significantly enriched within screen hits. mTor, a key nutrient sensor and positive regulator of translation (Laplante and Sabatini, 2012), is the top hit in the screen (Table S1). Validation of the RNAi screen was carried out by independent single gene knockdowns, using siRNAs that differ in sequence from the shRNAs used in the original screen. Importantly, knockdown of individual genes involved in translation and growth leads to significant decreases in Chd1chr-EGFP reporter intensity within 2-3 days, and in some cases the effect is stronger than loss of individual chromatin regulators (Figure 1E). These results suggest that translation and growth positively regulate euchromatin in ES cells.

### Translation, mTor and Myc activities promote euchromatin in ES cells

We next sought to expand upon the RNAi screen results using small molecule inhibitors of several growth-associated processes (Figure 2A). We found that acute inhibition of protein synthesis, mTor activity or Myc/Max complex activity leads to a rapid decrease in Chd1chr-EGFP fluorescence within 3 hours, while the Hp1α-EGFP reporter and an EGFP-only control remain unaltered during this time frame (Figure 2B-D). There is partial recovery from this effect by 24h, possibly due to a cellular adaptation to a lower growth state (Fig. S2). In contrast, inhibition of RNA Pol I or Pol II activity has no discernible effect on Chd1chr-EGFP fluorescence until 24 hours of treatment (Figure S2A), suggesting that inhibition of translation or cellular growth has a more immediate impact on ES cell euchromatin than general transcriptional inhibition.

**Figure 2.**
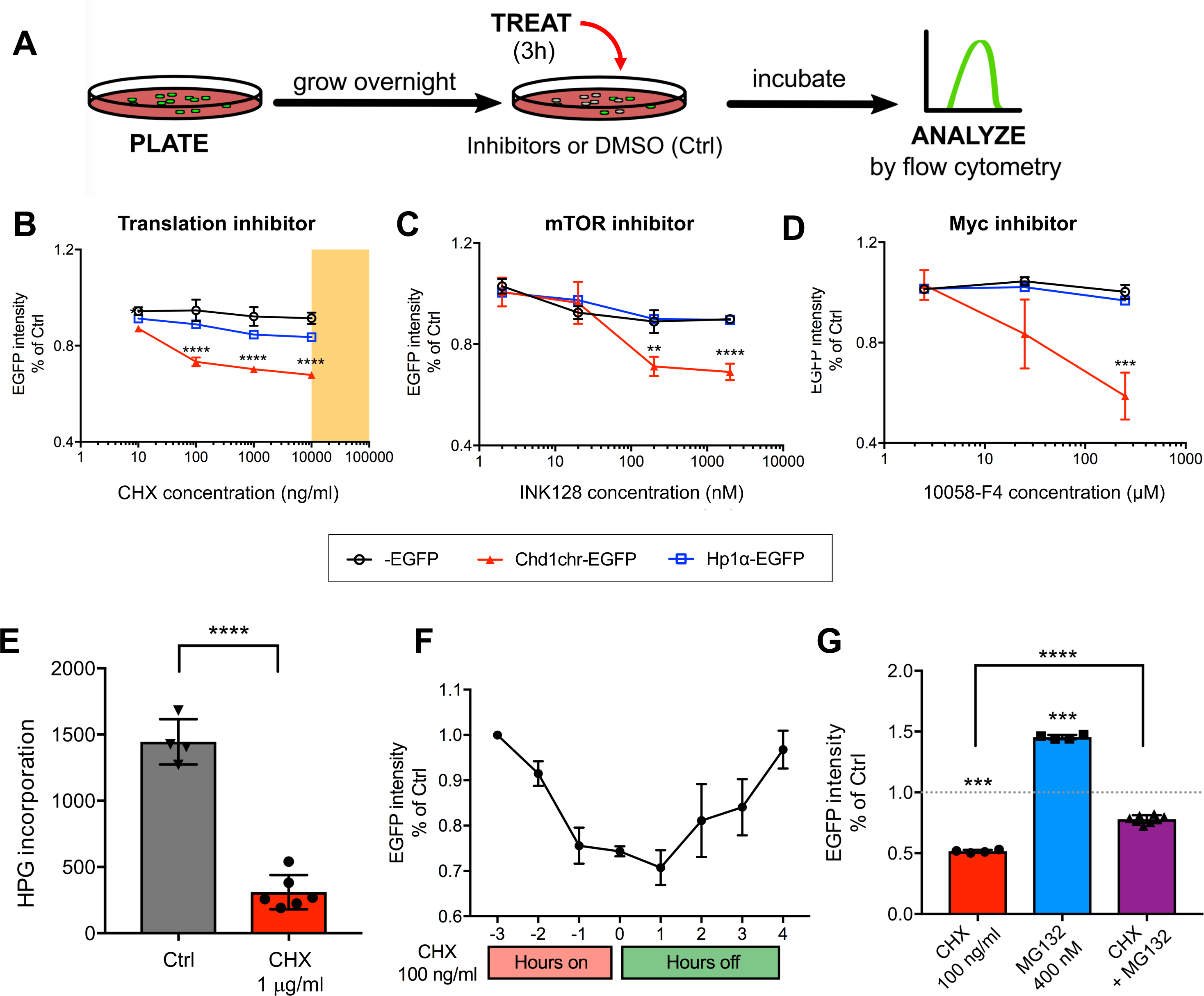
Translation, mTor and Myc dynamically regulate euchromatin reporter activity. (A) Schematic of small molecule-mediated inhibition and readout of selected pathways. See Fig S2 for other pathways tested. (B-D) Response of the Chd1chr-EGFP, Hp1α-EGFP and control EGFP ES cells to inhibition of translation (B), mTOR (C), or Myc/Max (D) at the indicated doses for 3 hours. Orange shading in (B) represents the range of CHX concentration used to fully inhibit translation. Cells were treated with DMSO as control. Graphs show mean ± SD of at least 3 technical replicates and are representative of 2 biological replicates. Statistical significance was determined by two-tailed t-test. **, ***, **** = p<0.01, 0.001, 0.0001. (E) Levels of nascent protein synthesis assessed by L-homopropargylglycine (HPG) incorporation in DMSO- or CHX-treated (3 hours) ES cells. Error bars show mean ± SD of at least 4 technical replicates and are representative of 2 experiments. (F) Recovery of the fluorescent chromatin reporter following CHX removal. CHX (at 100 ng/ml) or DMSO was added for 3 hours and then replaced with fresh medium for the next 4 hours. Graph depicts mean ± SD of 4 technical replicates and is representative of 2 biological replicates. (G) Proteasome inhibition partially rescues the effect of CHX on euchromatin. Reporter ES cells were treated with either DMSO, CHX, MG132 (proteasome inhibitor), or CHX and MG132 for 3 hours at the indicated doses. Fluorescence is reported as above.

ES cells cultured in Gsk3 and Mek/Erk inhibitors (2i condition) and LIF capture the naïve pluripotent state (Ying et al., 2008). 2i ES cells grow slower than ES cells in serum and LIF and have lower levels of global nascent transcription and steady-state RNAs (Bulut-Karslioglu et al., 2016). As expected, we found that 2i ES cells display significantly lower levels of Chd1chr-EGFP intensity (Figure S2B). Treatment of 2i ES cells with translation, mTor or Myc inhibitors further reduces Chd1chr-EGFP reporter activity (Figure S2C-E). These results support the notion that euchromatin rapidly adjusts to changes in growth and translational output of ES cells in different pluripotent states.

Cycloheximide (CHX) is normally used at 30-50 μg/ml to fully inhibit translation. Concentrations lower than 10 μg/ml enable translation to proceed, albeit with a reduced rate of elongation (Siegel and Sisler, 1963) (Figure 2E and S3A). Therefore, it is noteworthy that mild CHX treatment at 100- to 1000-fold lower concentrations than that required to completely block translation selectively decreases Chd1chr-EGFP intensity within 3 hours (Figure 2B), with no impact on cell survival (Figure S3B) and in a reversible manner (Figure 2F). Interestingly, inhibition of the proteasome partially rescues the decrease in Chd1chr-EGFP intensity induced by CHX, suggesting that protein turnover mediates the sensitivity of euchromatin to inhibition of translation (Figure 2G, and see below). Chd1chr-EGFP mRNA levels are stable and protein levels are only mildly affected after 3 hours of CHX treatment at 100 ng/ml (Figure S3C and D), suggesting that the decrease in fluorescence may be due to a delocalization of the fusion protein from the chromatin compartment upon alteration of the chromatin landscape. We focused on the acute impact of reduced translation on ES cell chromatin for the remainder of this study.

### Euchromatic histone marks are rapidly depleted upon inhibition of translation

We next explored the dependency of euchromatin on translation in a reporter-free system, using wild-type ES cells. Notably, inhibition of translation using CHX for 3h leads to a reduction in the levels of histone marks associated with active promoters and enhancers, such as H3K4me3 and H3/H4 acetylation, without affecting overall histone H3 levels or repressive H3K9me2 (Figure 3A). We confirmed the global reduction in acetylated H4 by intracellular flow cytometry (Figure S4A) and immunofluorescence assays (Figure 3B). Pluripotent cells in the ICM of the E4.5 blastocyst respond similarly to a mild 3h inhibition of translation, with rapid reductions in H4K16ac and H3K4me3 (Figure 3C and S4B). Serum starvation, which is known to reduce mTor activity and translational output, also leads to a decrease in H3K4me3 and H3K27ac within 3-6 hours (Figure S4C). Interestingly, H3K36me2 levels rise with increasing concentrations of CHX in a manner anti-correlated with H3/H4 acetylation (Figure 3A). These results agree with the observation that H3K36me2 recruits histone deacetylases to prevent spurious transcription (B. Li et al., 2009). Moreover, the levels of H3K9ac, which is deposited at TSSs and enhancers and correlates with H3K4me3, are distinct from other tested acetylation sites. H3K9ac accumulates in both ES cells and embryos upon CHX treatment at 100 ng/ml, yet it is depleted at higher concentrations (Figure 3A and S4B). Thus, deposition of H3K9ac may not be as strictly linked to H3K4me3 as previously thought (Karmodiya et al., 2012). Overall, these data suggest that the euchromatic compartment of ES cells is acutely sensitive to reductions in translational output.

**Figure 3.**
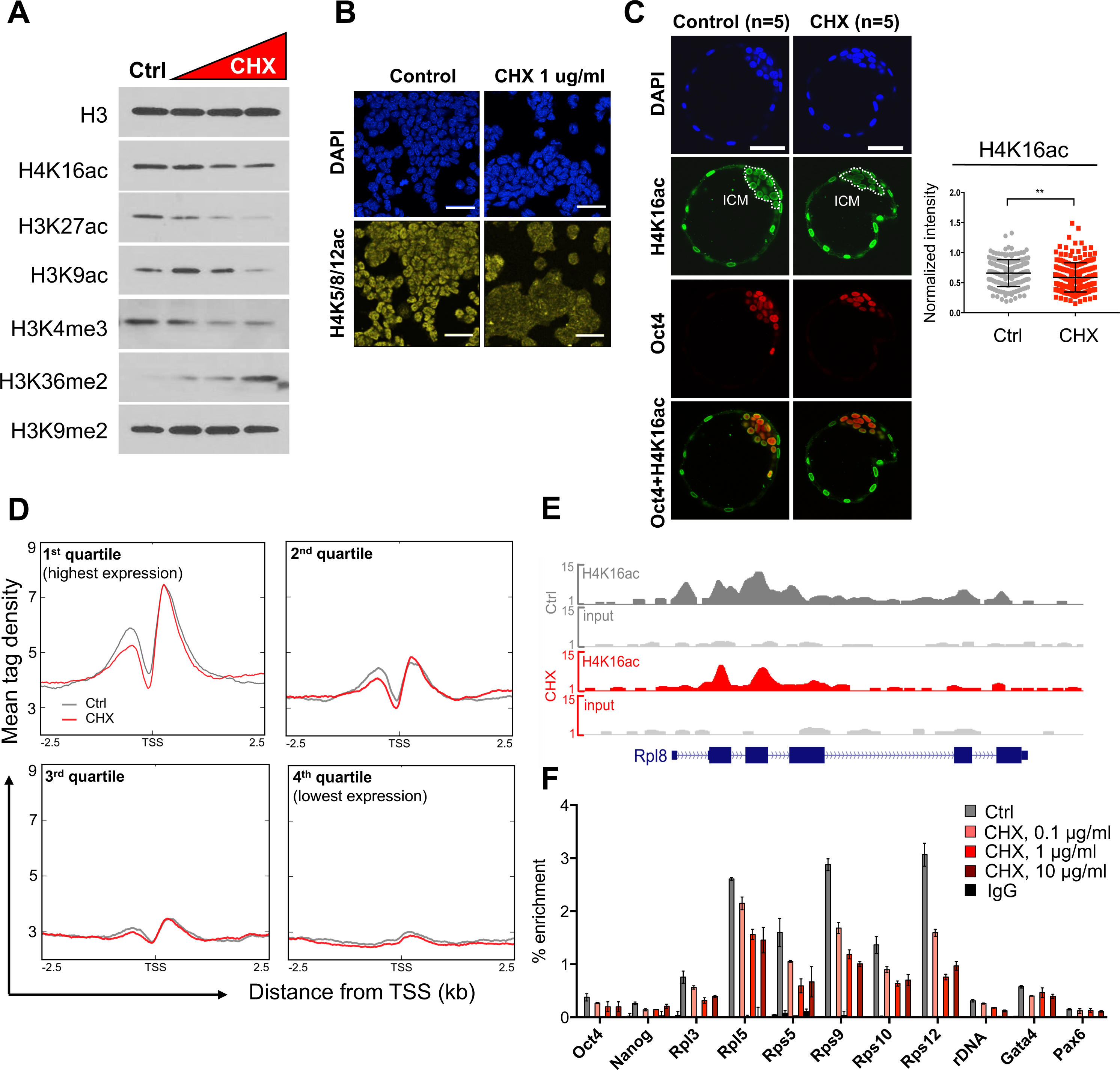
Inhibition of translation rapidly induces depletion of euchromatin marks in ES cells and blastocysts. (A) Levels of indicated histone modifications upon 3 hours of cycloheximide (CHX) treatment at 0.1, 1 or 10 μg/ml. Blots are representative of 2 biological replicates. (B) Immunofluorescent detection of H4 acetylation in control or CHX-treated ES cells. An antibody that recognizes acetylation on H4 K5, 8 and 12 was used. Scale bar denotes 50 μm. (C) Immunofluorescent detection of H4K16ac in control or CHX-treated (3 hours) E4.5 blastocysts. Blastocysts were flushed at E3.5 and cultured *ex vivo* until E4.5. DMSO or CHX was added in the last 3 hours of culture. A representative z-section of each embryo is shown. Scale bar denotes 50 μm. Right panel shows quantification of the H4K16ac signal in each Oct4+ cell. Statistical significance was determined by Welch’s two tailed T-test. ** = p<0.01. (D) Correlation of CHX-induced acetylation changes with gene expression level (CHX at 1 μg/ml for 3 hours). Genes were divided into 4 quartiles based on expression in ES cells (Bulut-Karslioglu et al., 2016). Profiles depict H4K16ac ChIP-seq tag density over annotated TSSs extended 2.5 kb upstream and downstream. (E) Representative genome browser view depicting H4K16ac in Ctrl or CHX-treated cells over the ribosomal protein gene *Rpl8*. (F) ChIP-qPCR documenting a dose-dependent response of H4K16ac following 3 hours of CHX. Error bars show mean ± SD of 3 technical replicates.

To gain insight into the genome-wide impact of inhibition of translation on histone acetylation, we performed ChIP-seq for H4K16ac after 3h of CHX treatment (Figure 3D-F, S4D-F). As previously described (X. Li et al., 2012), H4K16ac is concentrated around the TSSs of active genes and correlates well with expression levels (Figure 3D). We found that highly transcribed genes show the strongest reductions in H4K16ac levels, especially in the region immediately upstream of the TSS (Figure 3D and S4E). Ribosomal protein genes are particularly affected (Figure 3E and S4E). ChIP-qPCR confirmed these findings as well as the dose response of H4K16ac levels to the concentration of CHX, initially observed by western blot (Figure 3A, F). In contrast, H3K4me3 is only slightly reduced at TSSs upon CHX treatment (Figure S4F). Thus, H4K16 acetylation at highly transcribed genes is exquisitely tuned to the levels of translational output in ES cells.

### Translational output positively feeds back into nascent transcription

The results above led us to ask whether an acute inhibition of translation impacts nascent transcription. Remarkably, a 3h inhibition of translation results in a 50% decrease in global nascent transcription, assessed by measuring incorporation of 5-ethynyl uridine (EU) (Figure 4A). Transcription of both Pol II-transcribed mRNA and Pol I-transcribed rRNA transcripts is ~90% decreased upon treatment with CHX for 3h (Figure 4B). Steady-state levels of the same mRNA and rRNA transcripts remain relatively stable within this time frame (Figure 4C). Thus, inhibition of translation, a manipulation often used to study protein stability and turnover, has an unexpected and rapid impact on nascent transcription in ES cells.

**Figure 4.**
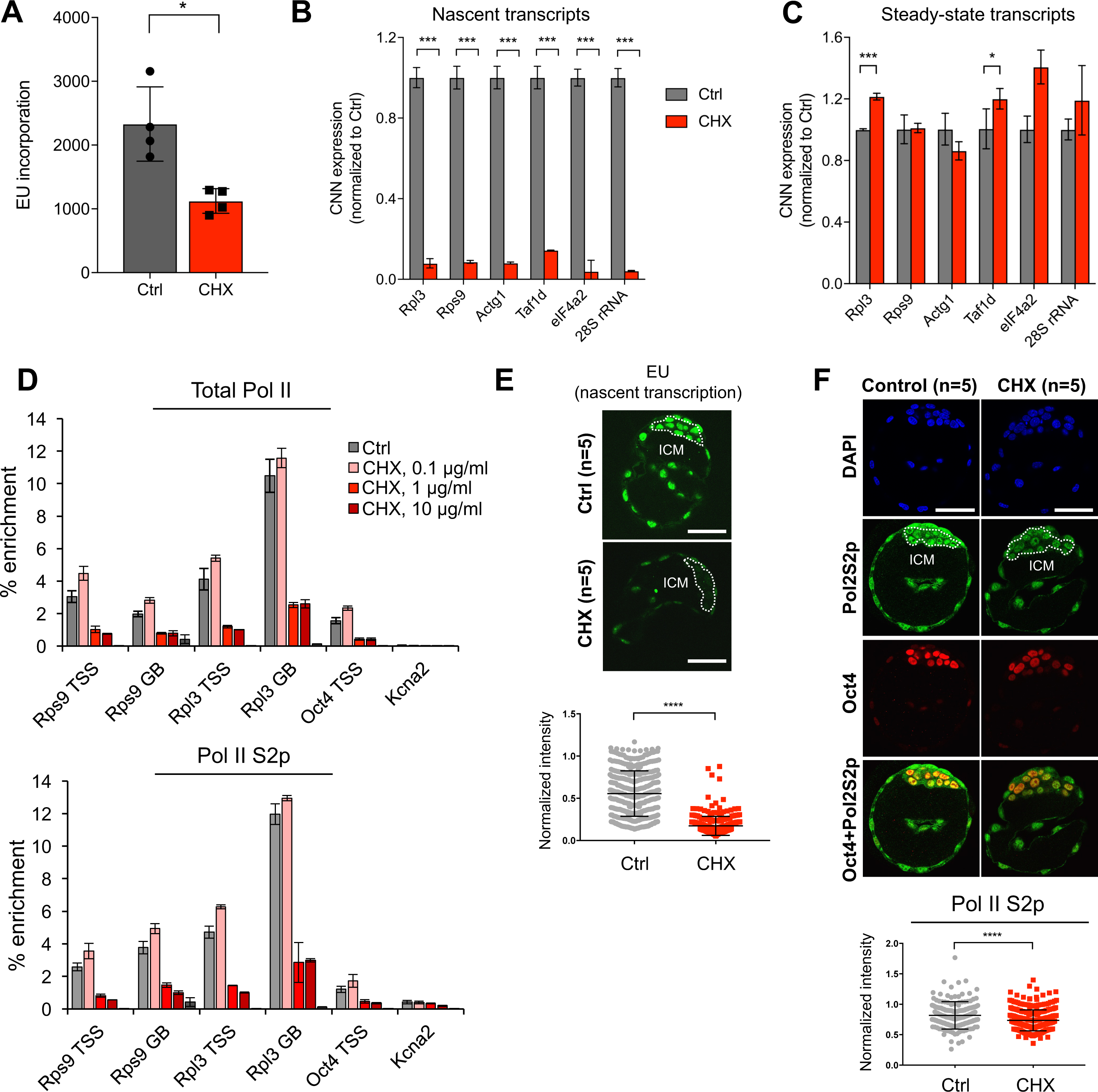
Nascent transcription is acutely sensitive to inhibition of translation in pluripotent cells. (A) Levels of global nascent RNA synthesis assessed by 5-ethynyl uridine (EU) incorporation in DMSO- or CHX-treated (3 hours) ES cells. (B) Nascent RNA capture followed by qRT-PCR in DMSO- or CHX-treated cells (see Experimental Procedures for details). Graph is representative of 3 biological replicates. Error bars show mean ± SD of 2-3 technical replicates. Statistical test performed was two-tailed t-test. *** = p<0.001. (C) Total mRNA levels of selected genes shown in (B) in DMSO- and CHX-treated cells. (D) Enrichment of total (upper panel) or elongating (lower panel) RNA Pol II at TSSs and gene bodies (GB) of selected genes in DMSO- or CHX-treated cells. Graph depicts show mean ± SD of 3 technical replicates. Graph is representative of 2 biological replicates. (E) Levels of nascent RNA synthesis assessed by EU incorporation in DMSO- or CHX-treated (3 hours) E4.5 blastocysts. Blastocysts were flushed at E3.5 and cultured ex vivo until E4.5. DMSO or CHX was added in the last 3 hours of culture and EU was added in the last 45 minutes of treatment. A representative z-section of each embryo is shown. Scale bar denotes 50 μm. Right panel shows quantification of the EU signal in the ICM (indicated by white dotted lines). Statistical significance was determined by Welch’s two tailed t-test. **** = p<0.0001.(F) Levels of elongating RNA Pol II (Pol2S2p) in control (DMSO) or CHX-treated (3 hours) E4.5 blastocysts. Bottom panel shows quantification of the Pol2S2p signal in each Oct4+ cell.

A reduction in nascent transcription could be due to increased pausing of RNA Pol II at the TSS or decreased occupancy at the TSS or along the gene body. ChIP-qPCR for total or elongating (Ser2p) RNA Pol II revealed that inhibition of translation leads to an overall decrease in polymerase occupancy at the TSS and gene body of highly expressed genes but not to increased promoter pausing (Figure 4D). We conclude that the decrease in nascent transcription is due to diminished recruitment or retention of RNA Pol II. Similar to ES cells, pluripotent cells of the blastocyst display significant decreases in nascent transcription (Figure 4E) and elongating RNA Pol II levels (Figure 4F) upon 3h inhibition of translation. Taken together, our data document that acute inhibition of translation not only alters the euchromatin landscape but also leads to a strong repression of transcription in pluripotent cells.

### Several key euchromatin regulators are unstable proteins

The sensitivity of euchromatin to reductions in translational output in pluripotent cells, and the partial rescue of Chd1chr-EGFP reporter levels observed upon proteasome inhibition (Figure 2G), led us to hypothesize that key euchromatin regulators may be unstable proteins that require continuous synthesis. To test this, we quantitatively assessed proteome-wide changes in ES cells after inhibition of translation using stable isotope labeling with amino acids in cell culture followed by mass spectrometry (SILAC-MS). In this case, we used a full block of translation (35 μg/mL of CHX) for either 1h or 3h to define the set of unstable proteins in ES cells (Figure 5A). We identified 4,078 unique proteins that were consistently depleted or enriched in the proteome at both time points of CHX treatment (Figure 5B and Table S3). Cell cycle factors are over-represented in the depleted proteins, a finding that is expected given that cell cycle progression is predominantly regulated by short-lived proteins (Table S4). Indeed, 3h inhibition of translation in ES cells moderately reduces the proportion of cells in S phase, with a concomitant increase in G_0_/G_1_ (Figure S5A, B). Fractionation of live cells in different stages of the cell cycle using a FUCCI reporter system (Nora et al., 2017) revealed that the impact of inhibition of translation on chromatin and transcription is observed throughout the cycle, although it is particularly evident in S/G_2_/M (Figure S5C-E). Regulators of chromatin, transcription and stem cell maintenance are also over-represented in the set of proteins rapidly depleted upon inhibition of translation (Figure 5B and Table S4). Overall, these results are in agreement with protein turnover data from *S. cerevisiae* and mouse fibroblasts, where cell cycle and transcription factors were found to be preferentially unstable (Belle et al., 2006; Schwanhäusser et al., 2011). Interestingly, a block in translation leads to a relative enrichment in the proteome of proteins associated with translation and mRNA/rRNA processing (Figure 5B). The translation machinery is generally stable, and is potentially least affected by changes in protein synthesis rate (Schwanhäusser et al., 2011).

**Figure 5.**
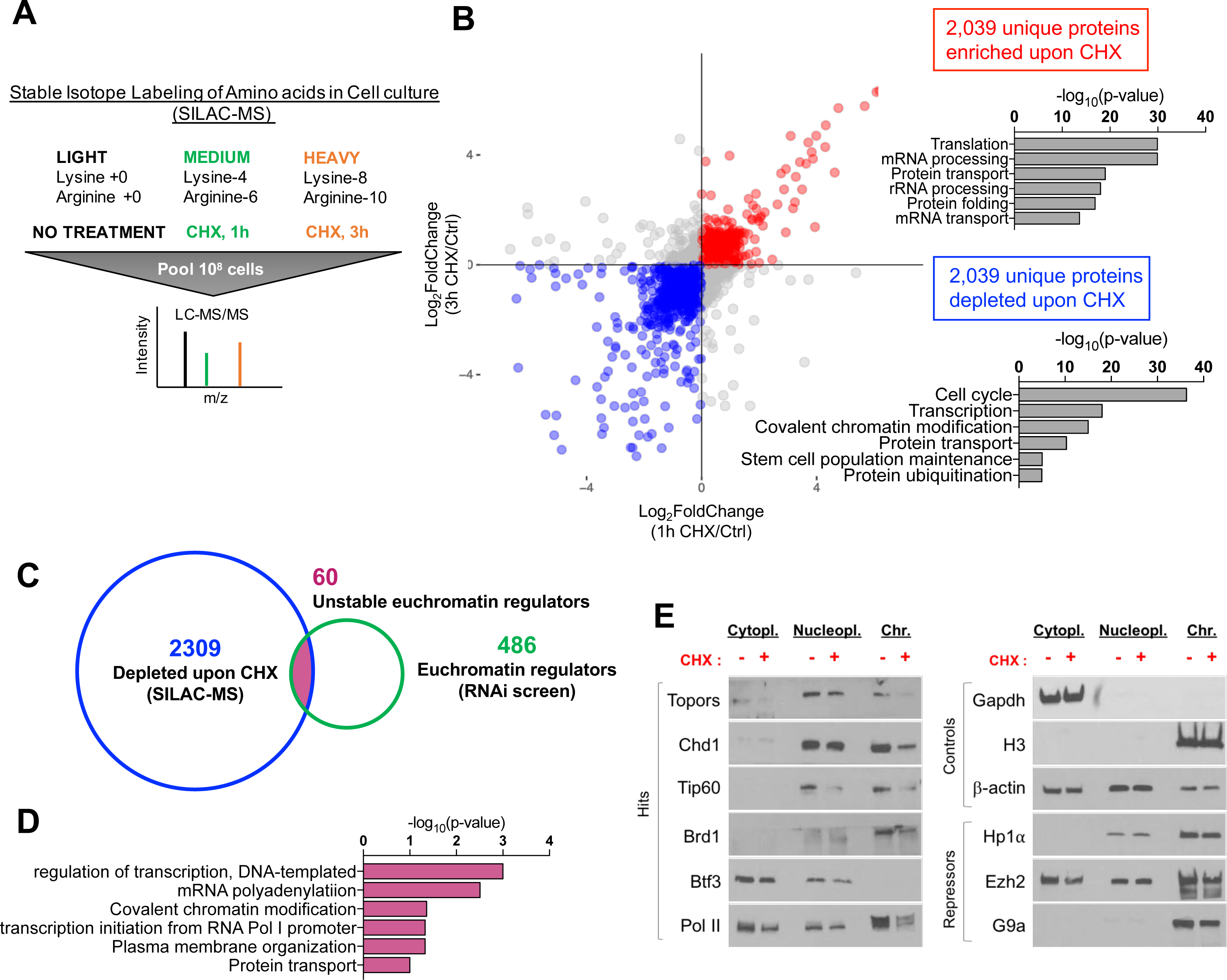
Key euchromatin regulators are unstable proteins that are rapidly depleted at the chromatin upon translation inhibition in ES cells. (A) Schematic of SILAC-MS workflow. (B) Scatter plot for proteins detected by SILAC-MS following 1 or 3 hours of CHX treatment. Proteins that are consistently enriched or depleted upon CHX treatment are shown in red or blue, respectively. Right panels show GO terms associated with the red and blue datasets. (C) Venn diagram for the intersection of RNAi screen hits with unstable proteins as determined by SILAC-MS (blue set in B). 60 such genes were identified. (D) GO terms associated with the 60 overlapping genes mentioned in (C). (E) Western blots showing the abundance of indicated proteins in cytoplasmic, nucleoplasmic and chromatin-bound compartments in DMSO- or CHX-treated (3 hours) cells. Left panel shows RNAi screen hits that are among the 60 proteins at the intersection shown in (C). Right panel shows proteins that serve as fractionation controls and chromatin regulators with repressor activities.

We next intersected the set of unstable proteins (Figure 5B, blue) with the RNAi screen hits (Figure 1B). This analysis yielded 60 proteins that are unstable euchromatin regulators (Figure 5C, D, Table S5). They include several known regulators of euchromatin and transcriptional activation in stem and progenitor cells, including Chd1 itself (Gaspar-Maia et al., 2009; Guzman-Ayala et al., 2014; Koh et al., 2015), the Tip60-p400 acetyltransferase complex (Fazzio et al., 2008), and the Brd1 component of the MOZ/MORF acetyltransferase complex (Mishima et al., 2011). We validated a representative subset as unstable proteins by western blotting after treatment with 3h CHX (Figure 5E). Interestingly, RNA Pol II is also selectively depleted from the chromatin fraction upon CHX treatment, in line with ChIP experiments (Figure 4D). In contrast, control proteins such as H3, Gapdh and β-actin, as well as the heterochromatin regulators Hp1α, Ezh2 and G9a, remain largely unchanged upon 3h CHX treatment. Thus, several key euchromatin regulators of ES cells are preferentially unstable proteins *in situ*, providing a mechanism for the acute dependence of permissive chromatin and hypertranscription on translation.

### A reduction in translational output rapidly deactivates developmental enhancers and primes transposable elements

Our studies to this point documented that a reduction in transcriptional output rapidly decreases the levels of activating histone marks and RNA Pol II at the promoters of highly expressed genes (Figures 3D-F, S4D, 4D), with a concomitant reduction in their nascent transcription (Figure 4B). However, the set of acutely unstable proteins in ES cells identified by SILAC-MS includes many sequence-specific transcription factors and chromatin regulators that are known to bind enhancers as well as promoters (e.g., Klf5, Gbx2, Zic1, Tip60/p400, Chd1, RNA Pol II, several Mediator subunits; see Figure 5 and Table S3). These results suggested that enhancer elements might also be sensitive to rapid shifts in translational output. We therefore carried out Assay for Transposase Accessible Chromatin with high throughput sequencing (ATAC-seq) to determine whether and how the landscape of chromatin accessibility in ES cells responds to acute (3h) inhibition of translation. We identified 454 regions that reproducibly lose accessibility and 734 regions that gain accessibility upon CHX treatment (Tables S6 and 7). Interestingly, most of these regions of differential accessibility are located 50-500 kb away from TSSs (Figure S6B).

Regions that become less accessible upon CHX treatment (CHX-lost) are associated with genes annotated with developmental functions (Figure 6A and Table S8). To probe the chromatin environment of CHX-lost regions, we analyzed published datasets (ENCODE Project Consortium, 2012; Goldberg et al., 2010; Ku et al., 2012). This analysis revealed that CHX-lost regions are active enhancers in wild-type ES cells (Calo and Wysocka, 2013), given their enrichment for DNase-hypersensitivity, H3K4me1, H3K27ac and p300 (Figure 6B). Thus, a subset of active enhancers associated with developmental functions in ES cells loses accessibility upon acute reduction in translational output. CHX-lost regions are highly enriched for DNA binding motifs of the transcription factors Klf4/5 and Zic1/3 (Figure 6B). Among those, Klf5 and Zic1 were detected at the protein level by SILAC-MS and are depleted upon 1h and 3h CHX treatment (Table S3). Klf4 has previously been shown to be an unstable protein (Chen et al., 2005). Therefore, reduced accessibility at active developmental enhancers upon inhibition of translation may be due to turnover of both euchromatin regulators and specific transcription factors with functions during development.

**Figure 6.**
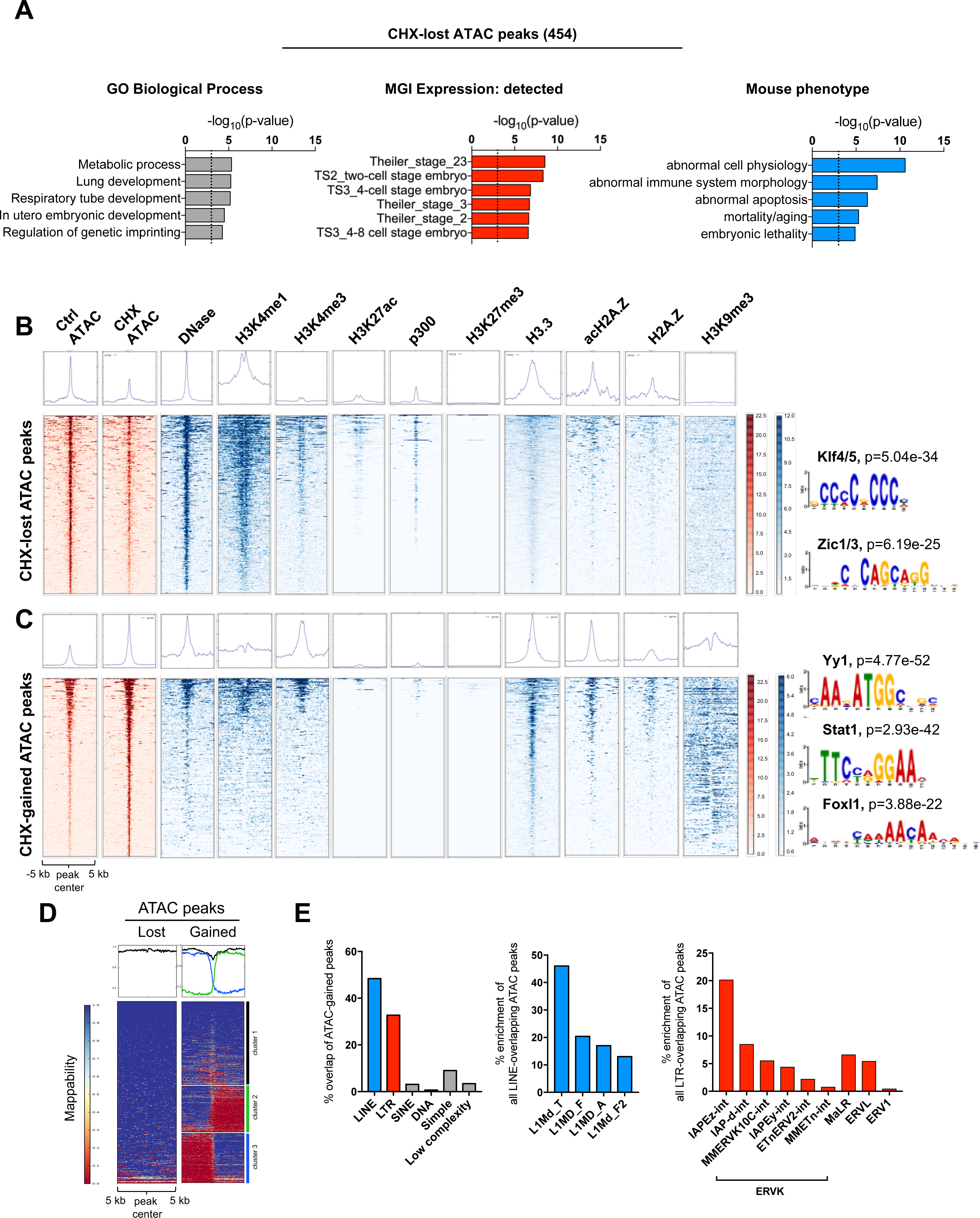
Inhibition of translation in ES cells induces reprogramming of chromatin accessibility at developmental enhancers, histone genes and transposable elements. (A) Functional terms associated with regions with loss of chromatin accessibility, determined by ATAC-seq, upon CHX treatment for 3 hours. See Table S7 for the full list of terms. (B, C) Heatmaps for enrichment of shown histone modifications, variants, and DNase-accessible sites on CHX-lost (B) or CHX-gained (C) ATAC-seq peaks. See Experimental Procedures for details on public datasets used. Right panels show DNA motifs highly enriched within CHX-lost or CHX-gained peaks. See supplementary files 1 and 2 for full lists of motifs. (E) Heatmaps showing levels of mappability of CHX-lost or CHX-gained ATAC-seq peaks. The CHX-gained heatmap is divided into three clusters to denote regions of distinct mappability. (F) Enrichment of repetitive elements over CHX-gained ATAC-seq peaks.

Regions that become more accessible upon CHX treatment (CHX-gained) are generally not associated with any gene or functional signature, with the exception of histone clusters (Figure S6C and Table S8). There is no significant accumulation of activating histone modifications on the majority of CHX-gained peaks. Rather, CHX-gained regions are embedded in domains of high levels of H3K9me3 (Figure 6C). We speculated that these regions might overlap genomic repeats. Indeed, CHX-gained peaks reside immediately upstream of long (~5kb) unmappable regions (Figure 6D). Moreover, ~50% and ~30% of CHX-gained peaks overlap with transposable elements (TEs) of the LINE1 and LTR families, respectively, and active subfamilies such as L1Md_T/F/A and IAP are particularly enriched (Figure 6E). Despite increased chromatin accessibility, the nascent expression of histone clusters and TEs is still suppressed upon CHX treatment, albeit to a lesser extent than mRNAs from non-repetitive genes (Figure S6D and E). The gains in chromatin accessibility upon acute inhibition of translation may be due to the fact that these regions are enriched for AA/AT dinucleotides, which tend to repel nucleosomes (Valouev et al., 2011), and are marked by H3.3 and (acetylated)H2A.Z, histone variants associated with nucleosome instability (Jin et al., 2009) (Figure 6C). Taken together, our results reveal that the open chromatin landscape of ES cells is rapidly reprogrammed upon partial inhibition of translation, with decreased accessibility at active enhancers associated with development and increased accessibility at histone genes and transposable elements.

## DISCUSSION

We report here that the transcriptionally permissive chromatin state of ES cells in vitro and ICM cells in vivo is acutely tuned to the levels of translational output. A genome-wide RNAi screen identified several novel regulators of euchromatin in ES cells, pointing to a central role for mTor and translation in this context. We found that a reduction in translational output very rapidly leads to global chromatin reorganization in ES cells, with preferential loss of histone marks associated with transcriptional activation and a global decrease in nascent transcription. An acute inhibition of translation further leads to loss of chromatin accessibility at a subset of developmental enhancers and increased accessibility at histone genes and repeat elements. Global analysis of protein stability in ES cells revealed that several key regulators of permissive chromatin are particularly unstable proteins and therefore require continuous synthesis. These findings point to a positive feedback loop between chromatin and translation, whereby the permissive, hypertranscribing chromatin state in ES cells not only promotes an elevated translational output but also depends directly on such elevated translation (Figure 7A). We propose that this feedback loop, in turn, sets the rapid pace of proliferation in ES cells and of embryonic growth at peri-implantation.

**Figure 7.**
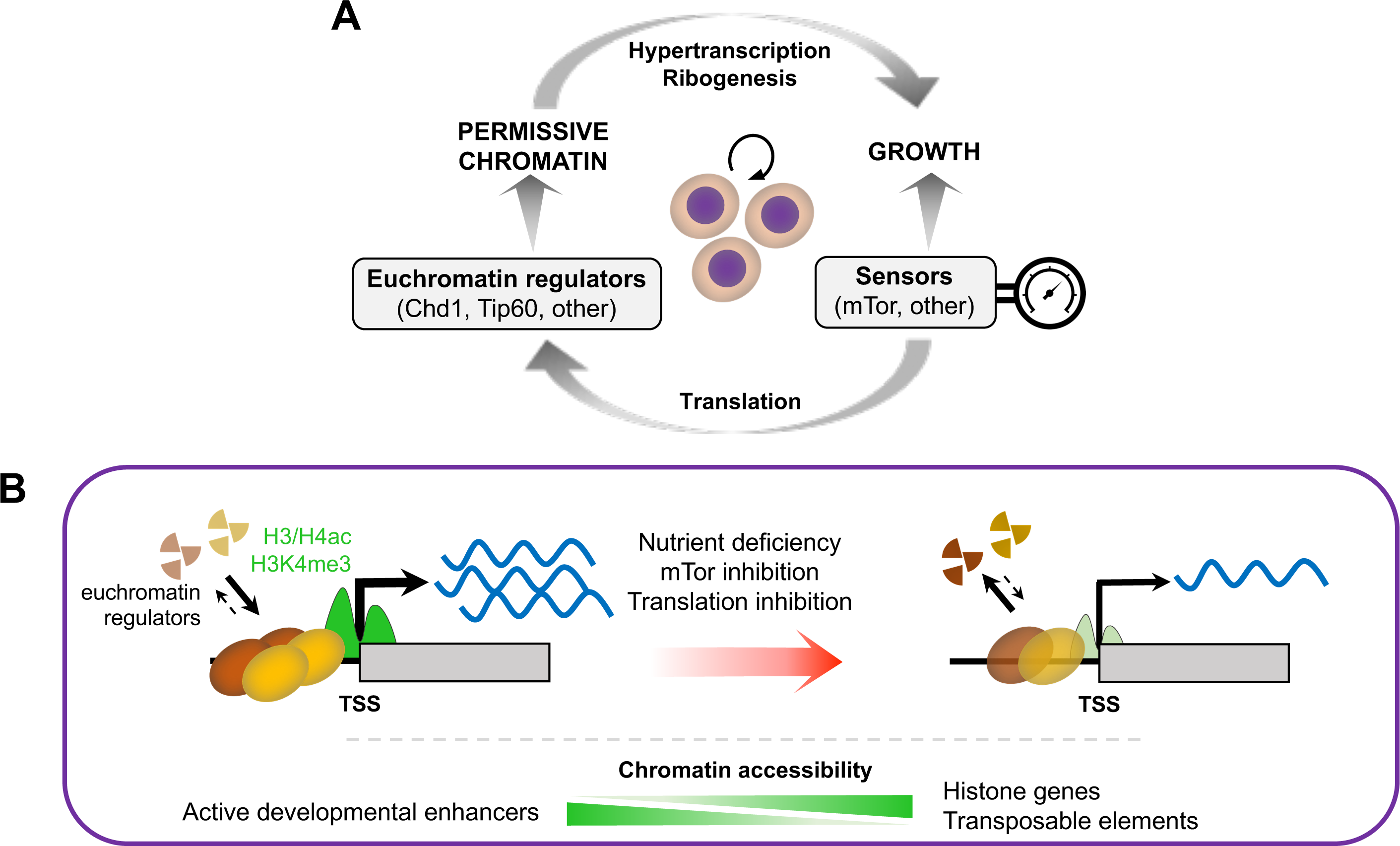
Proposed model for the dynamic feedback between translation, chromatin and transcription in ES cells. (A) The permissive chromatin state of ES cells promotes growth by sustaining hypertranscription and ribogenesis, whereas growth promotes the permissive chromatin state by sustaining high levels of translational output. Signaling and nutrient sensors such as mTor act as rheostats of this positive feedback loop. (B) The permissive chromatin state of ES cells responds rapidly to changes in translational output, in part due to the instability of euchromatin regulators. See Discussion for details.

Our results document a remarkably fast response of euchromatin and transcription to moderate perturbations of translation output in ES cells, whereas heterochromatin and its regulators appear to be more stable overall (Figure 7B). Histone acetylation may be a key integrator of the status of translation and nutrient availability in this context, given the instability of components of histone acetyltransferase complexes such as Tip60, p400 or Brd1 (Figure 5E) and the fact that histone acetylation is directly dependent on the glycolytic state of undifferentiated ES cells (Moussaieff et al., 2015). Moreover, histone acetylation controls the highly dynamic nature of euchromatin, but not heterochromatin, in ES cells (Melcer et al., 2012). The intricate relationship between different histone modifications associated with transcription, notably histone acetylation and H3K4me3 (Crump et al., 2011), likely contributes to propagate the impact of altered translational output across various layers of regulation of chromatin activity.

Considering the profound changes in levels of activating histone marks and nascent transcription after 3 hours of inhibition of translation, it is interesting that the overall landscape of chromatin accessibility at promoters and gene bodies is largely unaffected. These results suggest that, on a short time scale, nucleosome occupancy in these regions is relatively resistant to changes in chemical modifications of histones and RNA Pol II activity. At distal regions, acute inhibition of translation induces loss of chromatin accessibility at a subset of developmental enhancers and gain at repeats. The net effect may be to limit spurious differentiation and prime a return to the high level of expression of histone genes and repeat elements that is typical of proliferating ES cells (Efroni et al., 2008), once translational output is re-established. In addition, it is possible that reduced translational output primes the chromatin of conserved LINE1 and LTR elements for retrotransposition, potentially as a stress response (Fedoroff, 2012). It will be interesting to determine to what extent ES cells that recover from inhibition of translation are distinct from normally growing ES cells, including with regards to chromatin accessibility and TE activity.

While our findings document that protein instability is a key tuner of the euchromatic state of hypertranscription in ES cells; other layers of regulation such as RNA stability, signaling or metabolism, are expected to play important roles. In this regard, it is interesting that in mouse embryonic fibroblasts the set of factors found to be unstable at both the protein and mRNA level is enriched for chromatin/transcription functions (Schwanhäusser et al., 2011). It is also important to note that reductions in translational output can have multiple effects not limited to chromatin and transcription, notably on the cell cycle. We found that within 3 hours of partial inhibition of translation, the proportion of ES cells in S phase is reduced, with a concomitant increase in G_0_/G_1_ (Figure S5). Nevertheless, for the levels of CHX used here the proportion of ES cells in S phase remains high (40-60%) and, importantly, euchromatin marks and nascent transcription are reduced upon inhibition of translation in both populations of G_1_/S and S/G_2_/M cells (Figure S5D,E). Interestingly, histone acetylation is required for efficient activation of replication origins during S phase (Unnikrishnan et al., 2010), and loss of histone acetylation drives yeast in nutrient-limiting conditions to enter quiescence (McKnight et al., 2015). In addition, the major H4K16 acetyltransferase MOF directly binds to and maintains the expression of genes required for cell cycle progression in proliferating mouse embryonic fibroblasts (Sheikh et al., 2016). Such links between euchromatic histone marks, transcription, translation, glycolysis and cell cycle may serve to coordinate overall biosynthesis with rapid proliferation in ES cells in vitro and epiblast cells in vivo (Snow, 1977).

The highly dynamic levels of euchromatin regulators may be due both to the reported inefficiency of translation in ES cells (Sampath et al., 2008) and to control by the ubiquitination and sumoylation pathways (Buckley et al., 2012; Vilchez et al., 2012). Several proteins with roles in these pathways are hits in the RNAi screen (Table S1). For example, Usp9x is a deubiquitinase required for early development (Pantaleon et al., 2001) and self-renewal of neural stem/progenitor cells (Jolly et al., 2009). Topors is an E3 SUMO/Ubiquitin ligase that targets chromatin modifiers (Pungaliya et al., 2007) and is itself unstable in ES cells (Figure 5E). These and other proteins may help coordinate an euchromatic/transcriptional response to perturbations in translational output via modification of chromatin factors. It will be of interest to dissect the role of the instability of specific euchromatin regulators such as Tip60/p400 or Chd1, as well as the function of ubiquitination/sumoylation factors such as Usp9x or Topors, in maintaining the permissive chromatin state of ES cells.

The positive feedback loop between permissive chromatin and translational output identified here may drive rapid proliferation of undifferentiated pluripotent cells, but it cannot be perpetuated indefinitely. In this regard, it is noteworthy that mTor is the top hit in the RNAi screen (Table S1). mTor may directly regulate permissive chromatin and hypertranscription, given its role in promoting histone hyperactylation at the nucleolus and high levels of ribosomal RNA transcription (Tsang et al., 2003). Moreover, it was the identification of mTor as a key regulator of permissive chromatin in this study that led us to the finding that inhibition of mTor induces a reversible state of *hypo*transcription and developmental pausing in ES cells and blastocysts (Bulut-Karslioglu et al., 2016). The centrality of mTor in growth signaling, nutrient sensing, ribogenesis and translational regulation (Laplante and Sabatini, 2012) make it an ideal rheostat for the positive feedback between euchromatin and translation during development.

Beyond ES cells and early embryos, hypertranscription is employed by germline and somatic stem/progenitor cells during phases of growth and regeneration (Percharde et al., 2017a; 2017b). Recent studies have shown that rapidly expanding lineage-committed progenitors often have elevated levels of transcriptional and translational outputs relative to their parental stem cells (Blanco et al., 2016; Sanchez et al., 2016; Signer et al., 2014; Zhang et al., 2014). In contrast, a global reduction in translational output is characteristic of dormant states, such as developmental pausing (Bulut-Karslioglu et al., 2016; Scognamiglio et al., 2016) or hibernation (Frerichs et al., 1998) and can be induced by environmental stresses including nutrient deprivation, hypoxia, viral infection or exposure to toxins (Laplante and Sabatini, 2012; Olsnes, 1972; Toribio and Ventoso, 2010). We speculate that the acute dependence of euchromatin and transcription on translational output is a recurrent feature in stem/progenitor cells that is modulated by environmental perturbations.

## SUPPLEMENTAL INFORMATION

**Figure S1.**
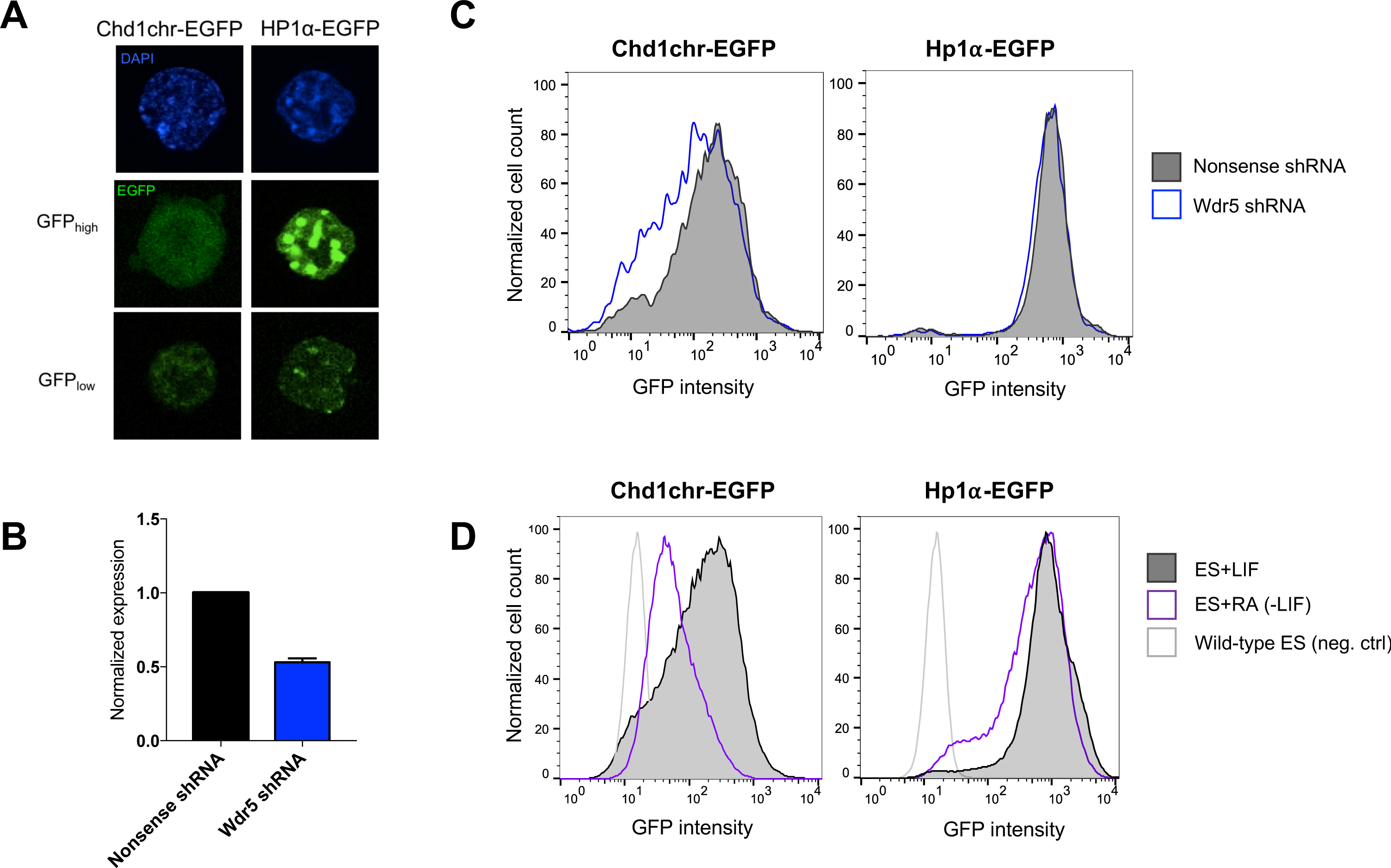
**Characterization of Chd1chr-EGFP and Hp1**α-**EGFP reporters in ES cells**. (A) Direct fluorescence imaging of the Chd1chr-EGFP and Hp1α-EGFP reporters. (B) mRNA expression levels of Wdr5, a component of the MLL1 complex that deposits H3K4 methylation, upon transduction of ES cells with non-targeting or Wdr5-specific shRNAs. (C) Flow cytometry analysis of Chd1chr-EGFP and Hp1α-EGFP reporter fluorescence levels upon knock-down of Wdr5. Fluorescence was assayed 3 days post-transduction. (D) Flow cytometry analysis of Chd1chr-EGFP and Hp1α-EGFP reporter fluorescence levels upon RA-mediated differentiation of ES cells for 2 days. ES cells grown in serum/LIF were used as control. Wild-type, non-fluorescent ES cells were used as negative control for flow cytometry.

**Figure S2.**
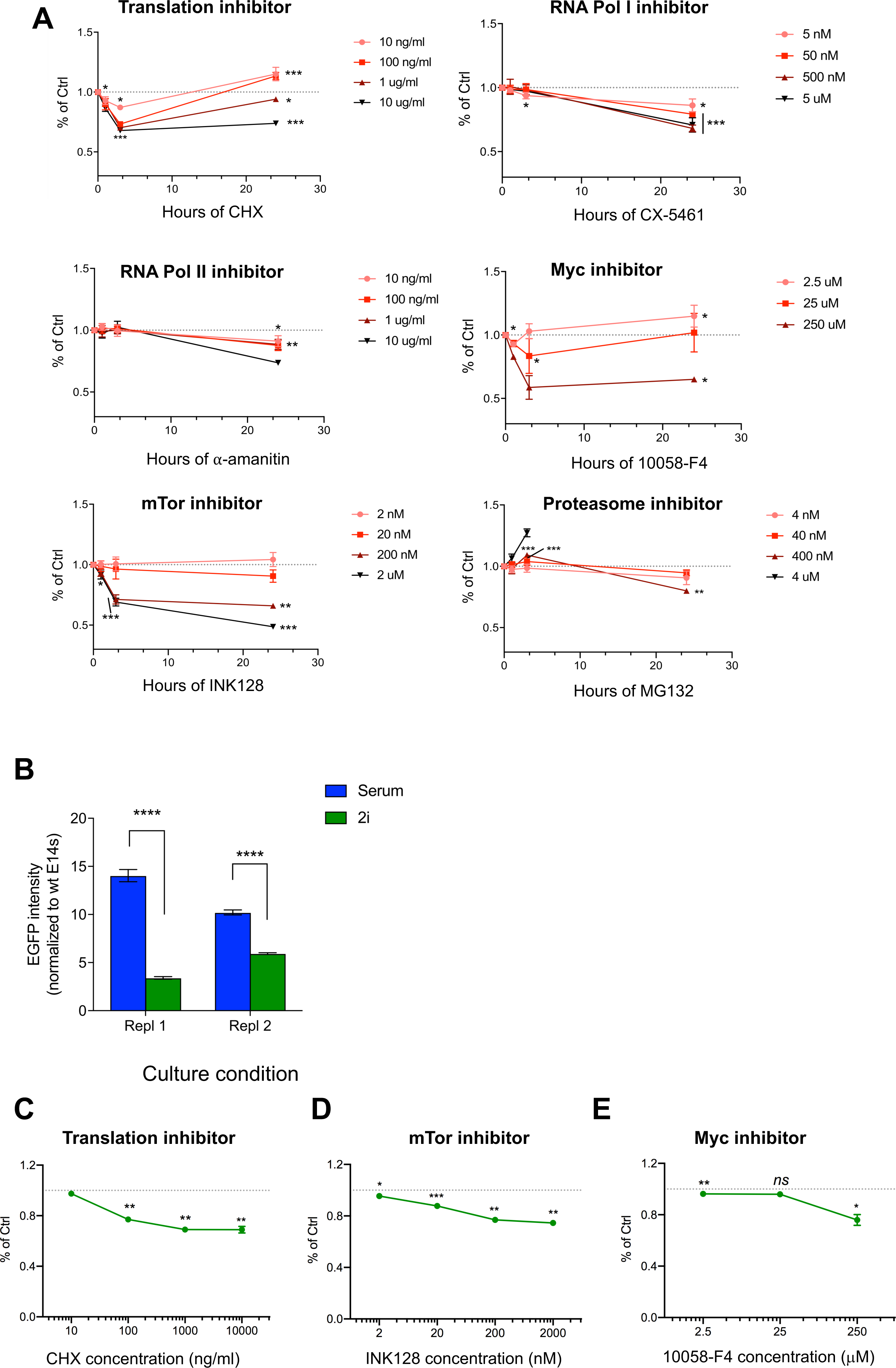
Characterization of the chromatin response to small molecule-mediated inhibition of indicated cellular pathways in 2i and serum ES cells. (A) Chd1chr-EGFP reporter fluorescence levels upon treatment with varying doses of the indicated inhibitors for up to 24 hours at indicated doses. Cells were treated with DMSO as control. (B) EGFP reporter expression in cells cultured in 2i or serum conditions. Fluorescence signal was normalized to wt (non-fluorescent) E14 cells. Error bars show mean ± SD of at least 8 technical replicates. Statistical significance was determined by Welch’s unpaired t-test. **** = p<0.0001. (C-E) Normalized fluorescence levels of the Chd1chr-EGFP reporter in 2i condition upon small molecule-mediated inhibition of indicated pathways for 3 hours. Data represent mean ± SD of 2 biological replicates. Statistical significance was assessed by Student’s unpaired *t-*test, assuming similar variances.

**Figure S3.**
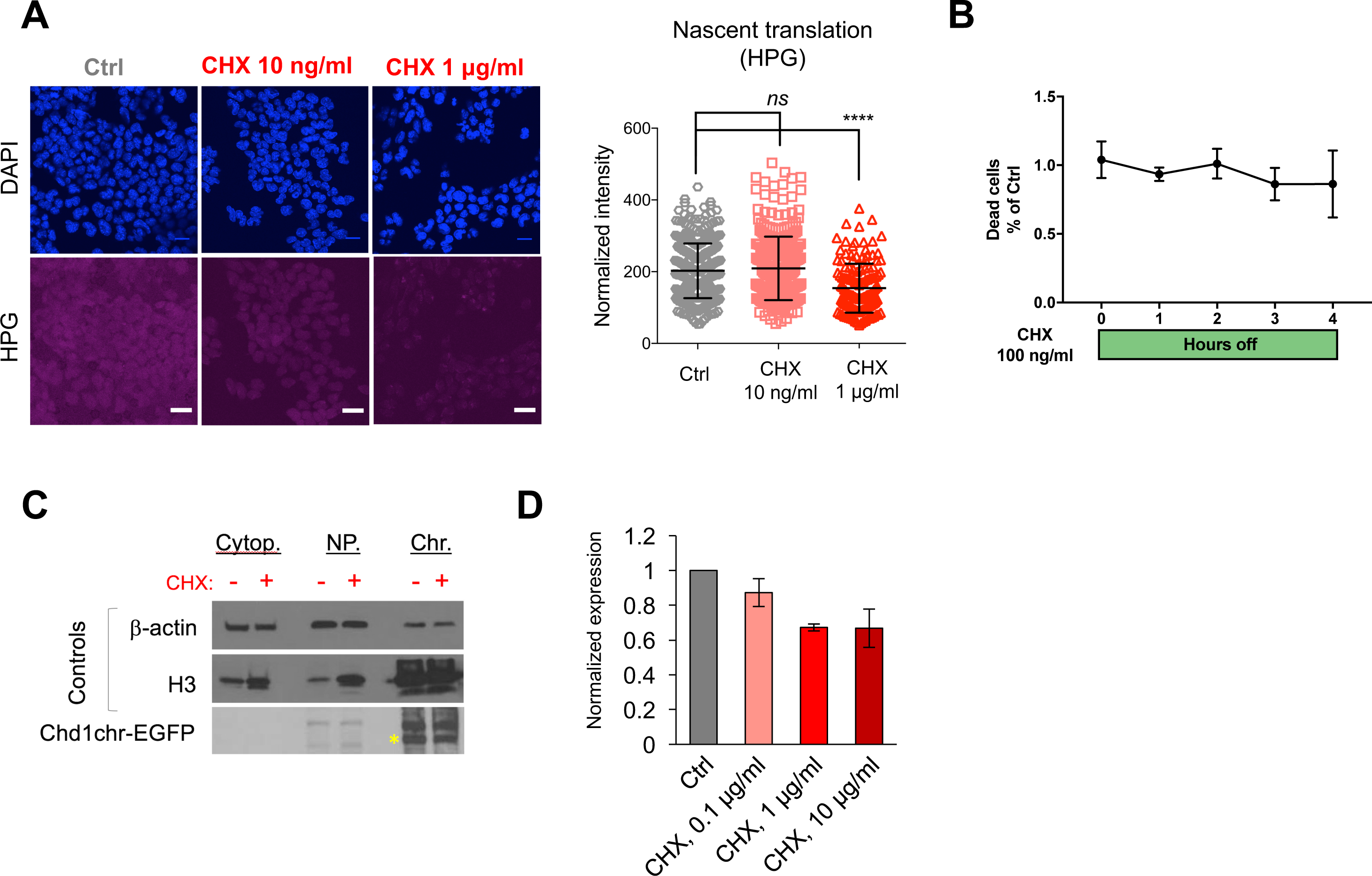
Effects of CHX treatment on global protein synthesis, cell survival and reporter expression. (A) Fluorescence imaging of nascent translation by HPG incorporation upon DMSO or CHX treatment. Scale bars represent 20 μm. Right panel shows quantification of HPG signal. Statistical analysis performed is Mann Whitney *U* test. Error bars represent mean ± SD of at least 3 technical replicates. (B) Assessment of cell death of CHX-treated ES cells by SYTOX Blue incorporation. Error bars show mean ± SD of 4 technical replicates. (C) Chd1chr-EGFP protein levels upon DMSO or CHX treatment for 3 hours. (D) Chd1chr-EGFP mRNA expression levels upon DMSO or CHX treatment for 3 hours. Error bars show mean ± SD of 3 technical replicates. Graph is representative of 2 biological replicates.

**Figure S4.**
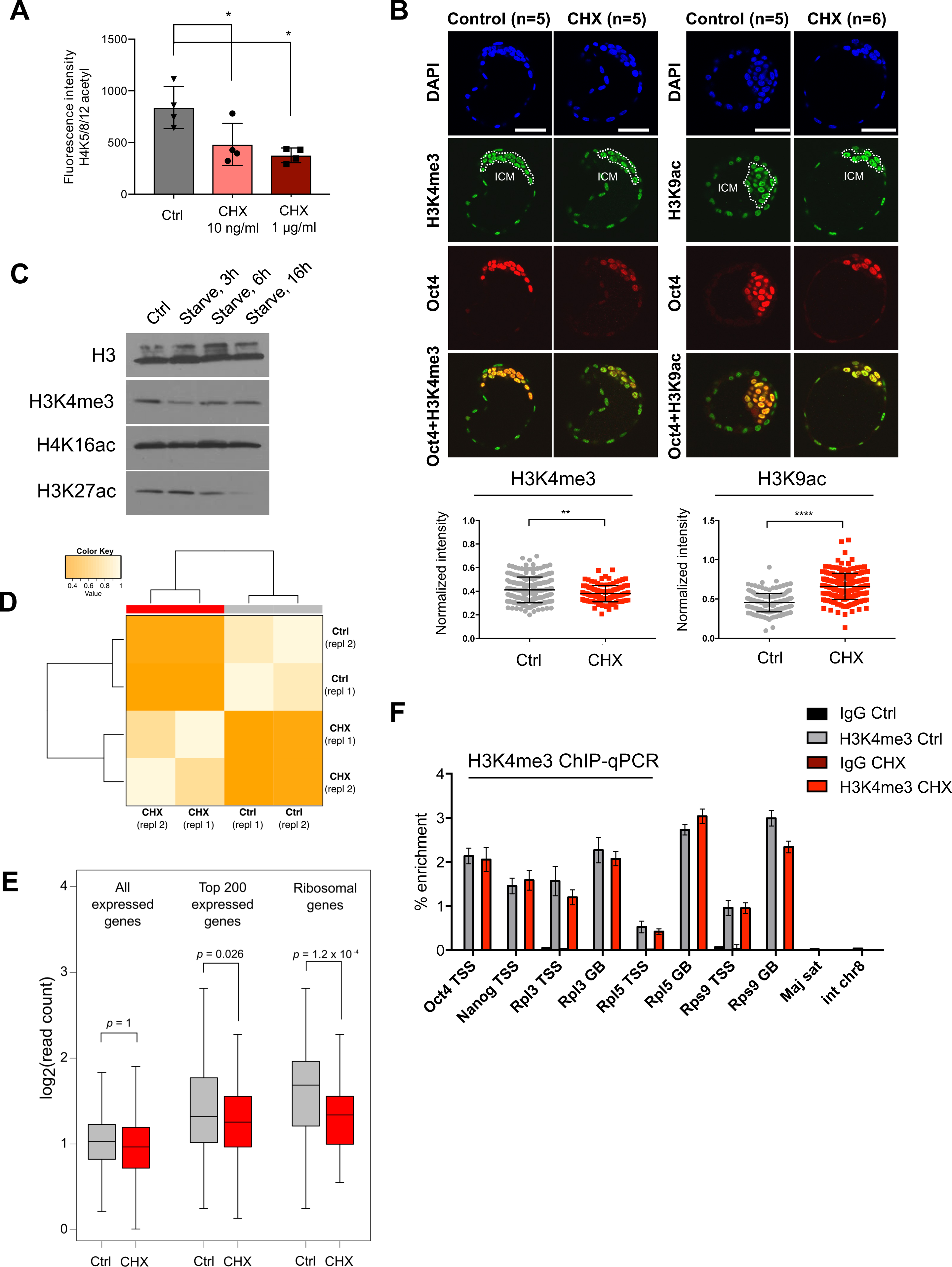
Chromatin response to inhibition of translation in ES cells and blastocysts. (A) Intracellular flow cytometry analysis of H4 acetylation (H4K5,8,12) in DMSO- or CHX-treated (3 hours) ES cells. Data shown are representative of 2 technical replicates. Statistical significance was determined by Mann Whitney *U* test. **** = p<0.0001. (B) Immunofluorescent detection of H3K4me3 and H3K9ac in control or CHX-treated (3 hours) E4.5 blastocysts. Blastocysts were flushed at E3.5 and cultured *ex vivo* until E4.5. DMSO or CHX was added in the last 3 hours of culture. A representative z-section of each embryo is shown. Scale bar denotes 50 μm. Bottom panels show quantification of the H3K4me3 or H3K9ac signal in each Oct4+ cell. Statistical significance was determined by Welch’s two tailed T-test. **, *** = p<0.01, 0.001. (C) Western blot analysis of euchromatin marks in response to serum starvation for the indicated durations. Histone extracts from unstarved cells were used as controls. Figure represents two biological replicates (D) Heatmap of H4K16ac ChIP-seq replicate correlation at the top 1000 most highly expressed genes in ES cells. (E) H4K16ac ChIP-seq read abundance over all expressed genes or gene subsets. (F) ChIP-qPCR for H3K4me3 enrichment over TSSs and gene bodies in DMSO-or CHX-treated cells (1 μg/ml, 3 hrs). Error bars show mean ± SD of 3 technical replicates.

**Figure S5.**
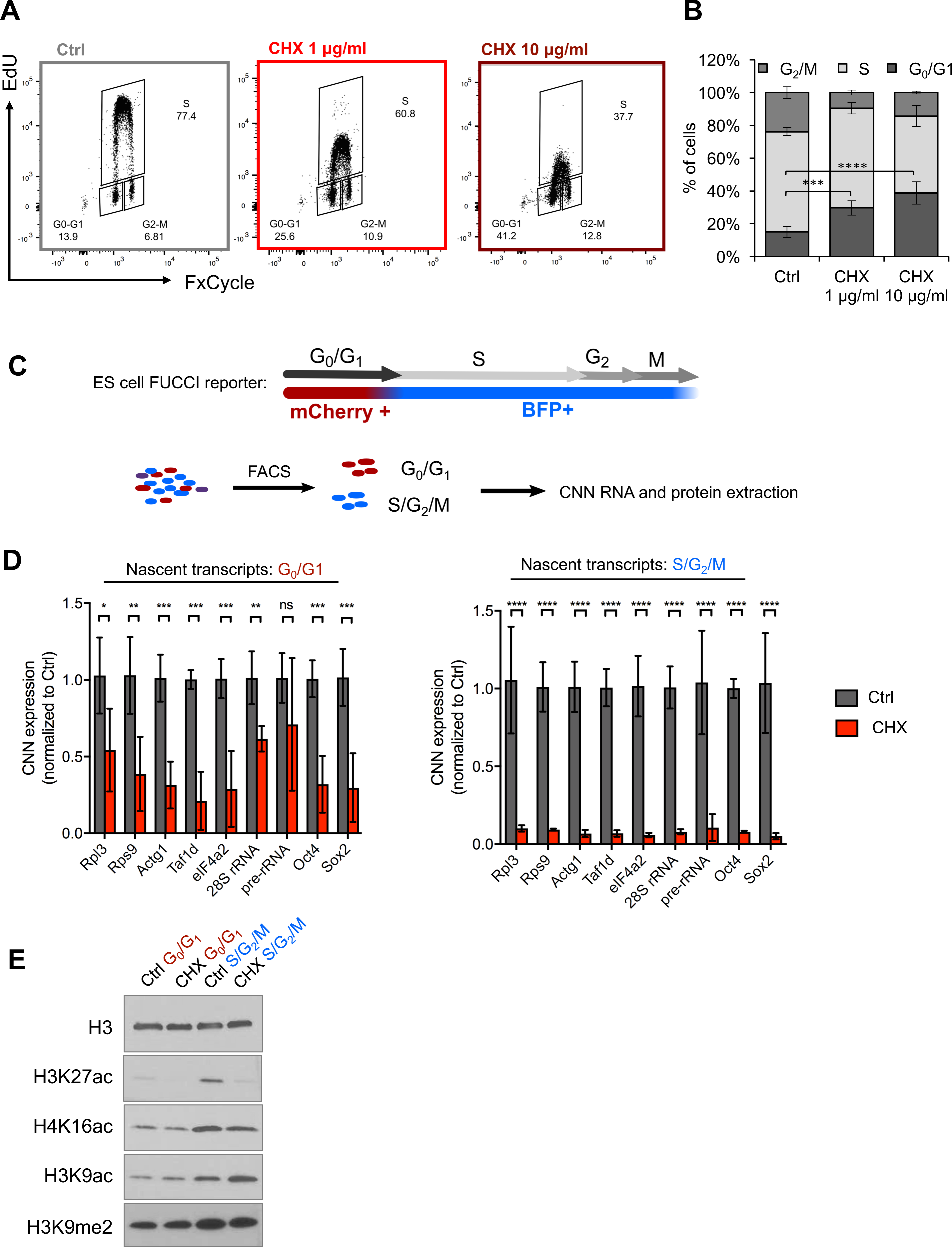
Impact of acute inhibition of translation on the cell cycle in ES cells. (A) Representative flow cytometry plots depicting cell cycle distributions of wild-type ES cells upon DMSO or CHX treatment. Cells were incubated with 1mM 5-ethynyl-deoxyuridine (EdU) for 1 hour. FxCycle staining was used to quantify DNA content. (B) Quantification of cell cycle stage distributions in DMSO- or CHX-treated ES cells. Error bars show mean ± SD of 2 biological replicates. Statistical significance was assessed by Student’s *t*-test. ***p < 0.001. (C) Schematics of the FUCCI cell line used in this study. (D) Nascent RNA capture followed by qRT-PCR in the indicated flow-sorted populations of DMSO- or CHX-treated (1 μg/ml, 3h) FUCCI cells (see Experimental Procedures for details). Error bars show mean ± SD of 2 biological replicates. Statistical test performed was two-tailed t-test. *** = p<0.001. (E) Levels of indicated histone modifications in flow-sorted populations of DMSO- or CHX-treated (1 μg/ml, 3h) FUCCI cells. Blots are representative of 2 biological replicates.

**Figure S6.**
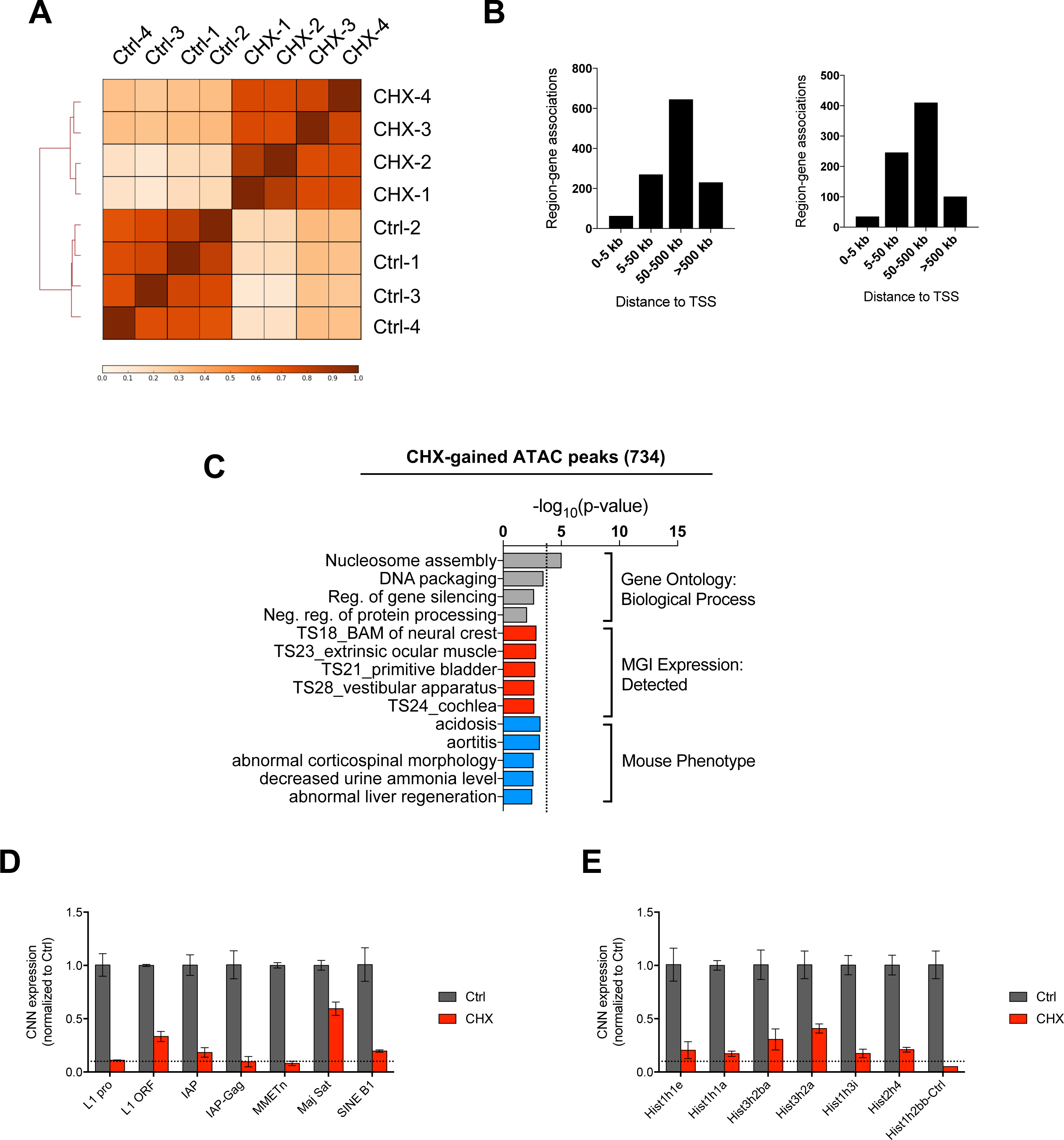
Characterization of chromatin accessibility and expression changes upon inhibition of translation in ES cells. (A) Unsupervised clustering of individual ATAC-seq replicates upon DMSO or CHX treatment for 3 hours. The top 10,787 most variable regions, as determined by Macs14 algorithm, were used for clustering analysis. (B) Distance of CHX-gained or CHX-lost regions from transcription start sites (TSS) (C) Functional annotation of ATAC-seq peaks lost upon CHX treatment for 3 hours. See Table S7 for the full list of terms. (D, E) Levels of nascent transcription of indicated transposable elements and histone genes in 3h DMSO- or CHX-treated ES cells, assessed by EU labeling followed by capture and qRT-PCR. Dotted lines represent the average level of downregulation for mRNAs depicted in Figure 4B.

**Table S1** RNAi screen hits

**Table S2** Gene ontology analysis of RNAi screen results

**Table S3** List of SILAC-MS-identified proteins

**Table S4** Gene ontology analysis of proteins depleted or enriched upon CHX treatment identified by SILAC-MS

**Table S5** List of 60 unstable euchromatin regulators

**Table S6** Coordinates of CHX-lost ATAC-seq peaks

**Table S7** Coordinates of CHX-gained ATAC-seq peaks

**Table S8** Gene ontology analysis of CHX-lost and CHX-gained regions identified by ATAC-seq

## ACKNOWLEDGMENTS

We thank Hiten Madhani and members of the Santos Lab for input and critical reading of the manuscript. We thank Thomas Jenuwein for the Hp1α-EGFP vector, Amanda Nolte for virus production, Richard Lao and members of the UCSF Institute of Human Genetics for assistance with sequencing and sonication, Shreya Chand for technical help, Ava Carter for advice on ATAC-seq libraries and Elphège Nora for sharing FUCCI ES cells. Samples were sequenced at UCSF Institute of Human Genetics Core Facility and Center for Advanced Technology, which is supported by the NIH (5P30CA082103). Flow cytometry data were generated in the UCSF Parnassus Flow Cytometry Core, which is supported by a Diabetes Research Center (DRC) grant and the NIH (P30 DK063720). SILAC-MS was performed at the UCSF Mass Spectrometry Facility, which is supported by the Biomedical Technology Research Centers program of the NIH National Institute of General Medical Sciences (NIH NIGMS 8P41GM103481 and NIH 1S10OD016229). This research was supported by NIH grant U01MH105028 and the UCSF Program for Breakthrough Biomedical Research (PBBR) to M.T.M., a Shurl and Kay Curci Foundation Research Grant and a gift from the Dabbiere family to A.D. and by NIH grants R01GM113014 and R01OD012204 to M.R.-S.

## AUTHOR CONTRIBUTIONS

A.B.-K. and M.R.-S. conceived of the project. A.B.-K. designed and performed the RNAi screen. A.D. analyzed the RNAi screen output. A.B.-K. and T.A.M. designed, performed and analyzed all other experiments except mass spectrometry. J.A.O.-P. performed mass spectrometry on SILAC samples. M.P. helped with bioinformatics analyses. S.C. and G.K. designed and cloned the shRNA library under the supervision of M.T.M. A.L.B. supervised the SILAC-MS experiment. M.R.-S. supervised the project. A.B.-K., T.A.M. and M.R.-S. wrote the manuscript with feedback from all authors.

## ACCESSION NUMBERS

Data have been deposited in Gene Expression Omnibus (GEO) under accession number GSE98358.

## EXPERIMENTAL PROCEDURES

### Mice

Swiss Webster females (6- to 12-week-old) and males (6 week- to 6 month-old) were used. Animals were maintained on 12 h light/dark cycle and provided with food and water *ad libitum* in individually ventilated units (Techniplast at TCP, Lab Products at UCSF) in the specific-pathogen free facilities at UCSF. All procedures involving animals were performed in compliance with the protocol approved by the IACUC at UCSF, as part of an AAALAC-accredited care and use program (protocol AN091331-03). Wild-type mice were mated, blastocysts were collected at E3.5 after detection of the copulatory plug by flushing uteri of pregnant females using M2 medium (Zenith Biotech) supplemented with 2% BSA (Sigma). Subsequent embryo culture was performed in 3.5-cm plates under light mineral oil (Zenith Biotech) in 5% O_2_, 5% CO_2_ at 37°C in KSOMAA Evolve medium (Zenith Biotech) with 2% BSA until E4.5.

### Generation of the Chd1chr-EGFP reporter cell line

A single PCR fragment containing the double chromodomains of Chd1was isolated and cloned in frame into the pCAGGS-EGFP-IRES-Puro plasmid containing a chicken β-actin promoter. Stable Chd1chr-EGFP expressing ES cell lines were generated by transfecting 4 μg *PvuI*-linearized vector into 10^6^ wt E14 ES cells, followed by Puromycin selection.

### ES cell culture

E14 (source: Bill Skarnes, Sanger Institute) ES cells were used for all reporter-free experiments. ES-FBS cells were cultured in DMEM GlutaMAX with Na Pyruvate (Thermo Fisher Scientific), 15% FBS (Atlanta Biologicals), 0.1 mM Non-essential amino acids, 50 U/ml Penicillin/Streptomycin (UCSF Cell Culture Facility), 0.1 mM EmbryoMax 2-Mercaptoethanol (Millipore) and 2000 U/ml ESGRO supplement (LIF, Millipore). ES-2i cells were cultured in DMEM/F-12, Neurobasal medium, 1x N2/B27 supplements (Thermo Fisher Scientific), 1 μM PD0325901, 3 μM CHIR99021 (Selleck Chemicals), 50 μM Ascorbic acid (Sigma) and 2000 U/ml ESGRO supplement (LIF) (Millipore). Cells were cultured in 0.5% FBS for serum starvation experiments. Cells tested negative for mycoplasma contamination.

### Wdr5 knock-down

Wdr5 shRNAs were designed based on the siRNA sequences from Ang et al (Ang et al., 2011). Control shRNA includes non-targeting sequence (Qin et al., 2014). shRNAs were cloned into the pSicoR-mCherry plasmid and constructs was packaged into lentivirus. Chd1chr-EGFP and Hp1α-EGFP ES cells were transduced with the lentiviral vectors. mCherry-positive cells with integrated shRNAs were sorted and knock-down was confirmed by qRT-PCR 72h post-transduction. EGFP fluorescence levels were analyzed on day 3 on an Avalon S3 Cell Sorter (Propel Labs).

### Retinoic-acid mediated differentiation

Chd1chr-EGFP and Hp1α-EGFP ES cells were used. LIF was withdrawn from ES-FBS medium and retinoic acid was added at a final concentration of 5 μM. EGFP fluorescence levels were analyzed on day 3 on an Avalon S3 Cell Sorter (Propel Labs).

### Western blot analysis

Histone analysis - Histones were acid extracted and TCA-precipitated as follows: 7×10^6^ cells were washed in ice-cold PBS with 5mM Na Butyrate. Cells were lysed in Triton Extraction Buffer (PBS,0.5% Triton X-100, 1× Protease Inhibitor Cocktail, 1 mM PMSF, 5 mM NaVO4 and 5 mM NaF) for 10 minutes and centrifuged. The pellet was resuspended in 0.2N HCl and histones were extracted overnight. Extracted histones were precipitated with TCA, washed with ice-cold acetone and resuspended in water.

Analysis of cellular fractions - Cytoplasmic, nucleoplasmic and chromatin fractions were isolated as previously described (Méndez and Stillman, 2000). Extracts were loaded into 4-15% Mini-Protean TGX SDS Page gels (Bio-Rad). Proteins were transferred to PVDF membranes. Membranes were blocked in 5% milk/PBS-T buffer for 30 min and incubated either overnight at 4°C or 1 hour at room temperature with the following antibodies: H4K16ac, H3K4me3, H3K9me3, Hp1α, Gapdh (Millipore), H3K27ac and H3K9ac (RevMAb Biosciences), H3, H3K36me2, H3K9me2, β-actin, RNA Pol II, RNA (Abcam), G9a, Ezh2 (Cell Signaling Technology), Topors, Chd1 (Santa Crux Biotechnology), Btf3 (R&D Systems), Brd1 (Developmental Studies Hybridoma Bank), Tip60 (GeneTex), EGFP (Thermo Fisher Scientific) and anti-rabbit/mouse/goat secondary antibodies. Membranes were incubated with ECL or ECL Plus reagents and exposed to X-ray films (Thermo Fisher Scientific).

For analysis of FUCCI ES cells, cells were plated overnight before 3h treatment with CHX at 1 μg/ml or DMSO. Cells were collected by trypsinization and sorted on a FACS AriaII (BD Biosciences) into mCherry+ (G_0_/G_1_) and BFP+ (S/G_2_/M) cell fractions. 4x10^5^ cells of each fraction were sorted for histone extraction and western blotting as above.

### Genome-wide shRNA screen

The ultracomplex EXPANDed shRNA library targeting the mouse genome was designed similarly to human shRNA libraries described before (Bassik et al., 2013; 2009). In brief, the library contains approximately 30 independent shRNAs per gene for all mouse protein-coding genes, for a total of ~600,000 shRNAs, hence the term “ultracomplex.“ The full list of shRNA sequences present in the library is available upon request. Pooled sequences coding for the shRNAs were cloned downstream of a U6 promoter in a modified pSicoR lentiviral vector containing a CMVPuro-T2A-mCherry cassette. All vectors were pooled to generate one lentiviral library representing the 600,000 shRNAs. The entire pooled library was then used to generate lentiviruses at the UCSF ViraCore. 6.6×10^7^ Chd1chr-EGFP ES cells were infected at an MOI (multiplicity of infection) ≤ 1 with the shRNA library, such that each shRNA is targeted to 100 cells (100× coverage). Cells were plated on thirteen 15 cm cell culture plates at a density of 5×10^6^ per 15 cm plate. Culture medium was changed daily; cells were harvested for analysis on day 3. mCherry-positive cells were sorted into GFP^low^ and GFPhigh populations on an Avalon S3 Cell Sorter (Propel Labs). Integrated shRNAs were isolated by PCR using oligos which contained sequencing adapters and barcodes. Screen results were analyzed as described before (Diaz et al., 2015).

### Single-gene knock down experiments

siRNAs were ordered as pools of 4 sequences from GE Dharmacon’s Cherry-Pick libraries. Chd1chr-EGFP ES cells were transfected with the siRNAs in 96-well plates. qRT-PCR and flow cytometry analyses were performed on day 2 or 3 as indicated. Fluorescence was analyzed on a BD Dual Fortessa.

### Inhibitor treatments

Chd1chr-EGFP ES cells were incubated for indicated durations and concentrations with the following inhibitors: INK128 (Medchem Express), 10058-F4 (Sigma), Cycloheximide (Amresco), α-amanitin (Sigma), CX-5461 (Selleckchem) and MG-132 (Selleckchem). Control cells were treated with DMSO. Inhibitors were withdrawn and cells were washed just prior to downstream analyses.

### Global nascent transcription and translation analysis

For measurements of global transcriptional and translational output, 4×10^5^ wild-type E14 cells were plated and cultured overnight. 5-ethynyl uridine (EU) was added at a final concentration of 10 mM, or L-homopropargylglycine (HPG) at a final concentration of 25 μM, and cells were incubated for the last 45 minutes of the 3 hour CHX treatment. Cells were trypsinized and counted (Bio-Rad Automated Cell Counter TC20, Bio-Rad). Equal cell numbers were collected from each sample and processed according to the instructions of the Click-iT RNA Alexa Fluor 488 HCS Assay kit (Thermo Fisher Scientific). Data were collected on a BD Dual Fortessa flow cytometer, analyzed using FlowJo v10, and plotted using Prism 7 (GraphPad). Datasets show similar variance.

### Nascent RNA capture followed by qRT-PCR

To measure nascent transcriptional changes at specific loci, ES cells were analyzed using the Click-iT Nascent RNA Capture Kit (Life Technologies). 4×10^5^ cells were plated and cultured overnight. The next day, 5-ethynyl uridine (EU) was added at a final concentration of 200 nM and cells were incubated for 30 minutes during DMSO or CHX treatment. Cells were washed, harvested by trypsinization and counted. Total RNA was isolated from the same number of DMSO- or CHX-treated cells using the Qiagen RNeasy Micro Kit (Qiagen) and processed according to manufacturer’s instructions. RNA was quantified using a Qubit 2.0 Fluorometer (Invitrogen). qPCR was performed with KAPA SYBR FAST qPCR Master Mix (Kapa Biosystems) and amplified on a 7900HT Real-time PCR machine (Applied Biosystems). Independent experiments were performed as above with 3 different cell numbers and RNA input amounts.

For analysis of FUCCI ES cells, cells were plated overnight before 3h treatment with CHX at 1 μg/ml or DMSO. EU was added at a final concentration of 200 nM for the last 30 minutes of drug treatment. Cells were collected by trypsinization and sorted on a FACS AriaII (BD Biosciences) into mCherry+ (G_0_/G_1_) and BFP+ (S/G_2_/M) cell fractions. 2×10^5^ cells of each fraction were sorted for RNA isolation and used for nascent capture as above.

### Immunofluorescent staining and imaging

Wild-type E14 ES cells were plated on gelatin in 8-chamber polystyrene vessels. Adhered cells were incubated with DMSO or CHX at the indicated concentrations for 3 hours. Cells were then fixed in 4% paraformaldehyde for 10 minutes, washed with DPBS and permeabilized with 0.2% Triton X-100 in PBS for 5 minutes. After blocking in PBS, 2.5% BSA, 5% donkey serum for 1 hour, cells were incubated overnight at 4°C with anti-H4K5/8/12 antibody (Upstate/Millipore). Cells were washed in PBS-Tween20, 2.5% BSA, incubated with fluorescence-conjugated secondary antibody (Life Technologies) for 2 hours at room temperature and mounted in VectaShield mounting medium with DAPI (Vector Laboratories). Imaging was performed using a Leica SP5 confocal microscope with automated tile scanning. Blastocyst staining and imaging was performed as described before (Bulut-Karslioglu et al., 2016) using blastocysts flushed at E3.5 and cultured until E4.5 (see Mice for details). CHX or DMSO was added in the last 3 hours of culture. All quantifications were performed using the Cell Profiler software (Carpenter et al., 2006).

### Intracellular flow cytometry

Wt E14 ES cells were cultured overnight in FBS/LIF before a 3-hour incubation in either CHX (100 ng/ml or 1 μg/ml) or DMSO (diluted to 1:10,000). Cells were fixed in 4% PFA for 15 minutes, permeabilized in 0.2% Triton X-100 for 3 minutes on ice, and blocked in 1% BSA in PBS. Primary incubation was performed with anti-H4K16ac antibody (Millipore 07-329) diluted 1:1000 in blocking solution, overnight at 4°C. Cells were washed and incubated in secondary antibody (AlexaFluor 488, Invitrogen) and fluorescence intensity was measured on a BD Dual Fortessa flow cytometer. Data were analyzed using FlowJo v10.

### H4K16ac ChIP-seq sample preparation

H4K16ac ChIPs were performed according to the recommendations of the Diagenode low-cell ChIP kit (Diagenode C01010070). Briefly, wt E14 cells were plated and cultured overnight. Cells were treated with CHX 1 μg/ml or DMSO diluted 1:10,000 in FBS/LIF medium for 3h. 10^5^ cells were harvested per IP. Lysis and IP were performed in the presence of 1× Halt Protease inhibitors (Thermo Scientific) and sodium butyrate (Sigma). Chromatin was sheared to an average size of 300 bp by a Covaris sonicator with the settings Duty 2, Intensity 3, 200 cycles per burst for 8 minutes. Shearing efficiency was checked by agarose gel. Fixed, sonicated chromatin was obtained from HEK 293 cells using the same method. IPs were performed using antibodies against H4K16ac (Millipore 07-329) or rabbit IgG (Abcam ab46540). Following overnight IP and washes, genomic DNA was treated with RNase A (Thermo Scientific). Reverse cross-linking was performed in the presence of Proteinase K (Clontech) at 65°C overnight. Genomic DNA was cleaned up by column purification (Qiagen Minelute Columns) and quantified by Qubit (Invitrogen). Two biological replicates were collected per condition.

### H4K16ac ChIP-Seq data analysis

HEK 293 chromatin was spiked in to a final concentration of 2.5% before library preparation. Sequencing libraries were prepared using the NEBNext ChIP-seq Library Prep for Illumina Kit (New England Biolabs) following manufacturer’s instructions. Libraries were constructed from 3 or 5 ng of DNA and quality was assessed by High Sensitivity DNA Assay on an Agilent 2100 Bioanalyzer (Agilent Technologies). Samples were sequenced on a HiSeq 4000 using single-end 50 bp reads. Sequencing reads that passed quality control were trimmed of adaptors using Trim Galore! and aligned to mm9 and hg19 using bowtie2 (Langmead and Salzberg, 2012) version 2.2.4 with no multimapping. Normalization factors for each sample, excluding inputs, were calculated from the ratio of total reads aligning to mm9 compared to total reads aligning to hg19 (Orlando et al., 2014). Reads were deduplicated using samtools (H. Li et al., 2009) and analyzed by custom R scripts, available upon request. Read coverage was assigned using the featureCounts function of the *Rsubread* package (version 1.24.1) (Liao et al., 2013) in R Bioconductor (Huber et al., 2015), using all coding genes of mm9 with a 2kb 5’ extension and disallowing multiple overlap. Read abundance over each gene was scaled by the respective normalization factor and then divided by the read abundance in the corresponding input. Replicate correlation was assessed by Pearson correlation between the top 1000 most-enriched genes, and replicates were pooled by summing the featureCounts of each replicate. Merged samples were used to produce all plots shown. Top-expressed genes were based on published CNN RNA-seq of E14 ES cells (Bulut-Karslioglu et al., 2016). Tag density plots were produced using deepTools on Galaxy (http://deeptools.ie-freiburg.mpg.de/) (Afgan et al., 2016; Ramirez et al., 2016). Box plots were produced using custom R scripts. Tracks were visualized using the UCSC Genome Browser.

### RNA Pol II ChIP-qPCR

ChIP was performed as described before (Brookes et al., 2012) using total RNA Pol II and RNA Pol II S2p (Abcam) antibodies and Protein A Dynabeads (Thermo Fisher Scientific).

### SILAC-Mass Spectrometry

To differentially label wild-type E14 ES cells with light, medium and heavy amino acids, we replaced the following components in the ES-FBS culture medium (see above): DMEM formulated without lysine and arginine instead of DMEM, dialyzed serum instead of regular FBS, lysine and arginine added separately to light (regular L-lysine and L-arginine), medium (L-lysine 4,4,5,5-D4 and L-arginine 13C6) and heavy (L-lysine 13C6, 15N2 and L-arginine 13C6, 15N4) media. All SILAC reagents were purchased from Cambridge Isotope Laboratories. L-proline was included in all media (included in with other non-essential amino acids) and the absence of arginine-to-proline conversion was verified by MS. Complete labeling (>99%) was confirmed by MS before starting the experiment. To quantitatively identify changes in protein levels upon inhibition of protein synthesis, cells were treated with DMSO (light) or 35 μg/ml CHX for 1h (medium) or 3h (heavy). Cells were washed with PBS and lifted in ice-cold PBS with 1× protease inhibitor cocktail (Roche). 10^8^ cells from each condition were pelleted and snap-frozen until MS analysis.

Harvested cells were lysed in 8M urea in 80 mM NH_4_HCO_3_, sonicated on ice (3 pulses at 35% power, 20 s each) and centrifuged at 15000g for 10 min. Supernatant was collected and protein concentration estimated with BCA. Equal amounts (100 ug) of the 3 SILAC labeled samples were combined, and the proteins treated with 5 mM DTT at 56C for 10 min, and then with 10 mM iodoacetic acid at RT for 1 h, then diluted 4 times to 2 M urea and digested o/n with trypsin (2% of total protein) at 37C. Samples were acidified with 5% formic, and peptides were extracted using SepPack cartridges. 200 ug of tryptic peptides where resuspended in 20 mM ammonium formiate pH 10.3, and separated in a 1 × 100mm Gemini 3um C18 column (Phenomenex) in a MeCN gradient (2 to 30 % in 60 min) in the presence of 20 mM ammonium formiate pH 10.3. 70 fraction were collected and combined in 18 final fractions, evaporated, resuspended in 0.1 formic acid and analyzed by LCMSMS in a Q Exactive Plus mass spectrometer (Thermo Fisher Scientific, USA). Peptides were loaded in a 200 cm monolithic C18 silica column (GL Sciences, Tokyo, Japan), and separated in a gradient of acetonitrile (288min 2 to 25%, 36 min to 32%, 18 min to 40%, 18 min to 60%, 5 min to 8%) in 0.1% formic The liquid chromatography eluate was interfaced with a 7 ◽m ID EasySpray emitter (Thermo Scientific) to the MS. Samples were analyzed in positive ion mode, and in information-dependent acquisition mode to automatically switch between MS and MS/MS acquisition. MS spectra were acquired in profile mode in the m/z range between 350-1500 m/z at 70,000 resolution. All samples were analyzed with a TOP10 method, the 10 most intense multiple charged ions over a threshold of 17000 counts were selected to perform HCD experiments. Product ions were analyzed in centroid mode with resolution R=17500, and isolation window was set to 4 Th. A dynamic exclusion window was applied that prevented the same m/z from being selected for 10 s after its acquisition.

Peaklists were generated using PAVA in-house software {Guan:2011kp} based on the RawExtract script from Xcalibur v2.4 (Thermo Fisher Scientific, San Jose, CA). The peak lists were searched against the human subset of the UniProt database as of June 17, 2013, (73955/36042779 entries searched) using in-house ProteinProspector version 5.10.10 (a public version is available online) with settings described below. A randomized version of all entries was concatenated to the database for estimation of false discovery rates in the searches. Peptide tolerance in searches was 20 ppm for precursor and 30 ppm for product ions, respectively. Peptides containing two miscleavages were allowed. Carboxymethylation of cysteine was allowed as constant modification; acetylation of the N terminus of the protein, pyroglutamate formation from N terminal glutamine, oxidation of methionine, and loss of the protein initial methionine, were allowed as variable modifications, as well as 2H(4) labelling in lysine and 13C(6) labelling in arginine, and 15N(2) 13C(6) labelling in lysine and 15N(4) 13C(6) labelling in arginine, limiting the allowed combination of labels: if K and R occur in the same peptide, Protein Prospector only allows pairing of K0 with R0, K4 with R6 and K8 to R10. In all cases, the number of modification was limited to two per peptide. A minimal ProteinProspector protein score of 20, a peptide score of 15, a maximum expectation value of 0.05 and a minimal discriminant score threshold of 0.0 were used for initial identification criteria. FDR was limited to 1%.

Quantification: SILAC quantification measurements were extracted from the raw data by Search Compare in Protein Prospector (http://prospector.ucsf.edu). Search Compare averaged together MS scans from −10 s to +30 s from the time at which the MS/MS spectrum was acquired in order to produce measurements averaged over the elution of the peptide. SILAC ratios were calculated, and base 2 logarithms of these values were used for further analysis. If quantitative data are available from isotopic envelopes identified as different charge states of the same peptide, the median of the log2 of the calculated SILAC ratios was used for that peptide. For proteins, the median of all the log2 ratios for peptides unique to that protein was calculated, and the distribution of log2 ratios normalized by its median value.

### ATAC-seq protocol and data analysis

ATAC-seq was performed as described before (Buenrostro et al., 2001) on 50,000 cells each of DMSO- or 1 μg/ml CHX-treated E14 ES cells, in quadruplicates. A total of 11 cycles of amplification was performed. Library quality and quantity were analyzed by Bioanalyzer (Agilent) and KAPA library quantification kit for Illumina platforms (KAPA Biosciences). Samples were sequenced on a HiSeq 4000 using single-end 50 bp sequencing reads. Raw reads were trimmed of adaptors and aligned to the mouse genome build mm10 using Bowtie2 version 2.2.4. Reads were deduplicated, sorted, and converted to bigWig format using samtools. BigWig files from individual replicates were used to assess correlation strength between replicates (Figure S6) using deepTools/multibigwigsummary and plotCorrelation tools. Peak calling was performed on each biological replicate pair using Macs14 version 1.4.2 20120305. Peaks from different biological replicates were then intersected. Peaks which were detected in at least 3 out of 4 replicates were selected for further analysis. As such, 734 peaks were found to be gained and 454 peaks were found to be depleted upon CHX treatment. Gene ontology analysis of these peaks was performed using GREAT software (McLean et al., 2010). The region-gene association was confined to 5 kb upstream and 1 kb downstream of each gene. To compare enrichment of histone marks, variants and DNase-seq signal over our ATAC-seq peaks, we used the following datasets: ENCODE (ENCODE Project Consortium, 2012) H3K4me1 (GSM1003750), H3K4me3 (GSM1003756), H3K27ac (GSM1000126), H3K9me3 (GSM1003751), H3K27me3 (GSM1000089), DNase (GSM1014154), p300 (GSM918750), additionally H2A.Z (GSM958501) and acH2A.Z (GSM958502) (Ku et al., 2012), H3.3 (GSM423355) (Goldberg et al., 2010), mappability tracks (UCSC table browser (Karolchik et al., 2004)). All heatmaps were generated using the deepTools package (Ramirez et al., 2016) on the Galaxy platform (Afgan et al., 2016). Motif analysis was done using Homer version 4.7 (Heinz et al., 2010). Analysis of repeats was done using repeat annotations from UCSC, custom R scripts and Galaxy Bedtools (usegalaxy.org).

